# Fast spiking interneurons autonomously generate fast gamma oscillations in the medial entorhinal cortex with excitation strength tuning ING–PING transitions

**DOI:** 10.1101/2025.09.05.674527

**Authors:** Brandon Williams, Ananth Vedururu Srinivas, Roman Baravalle, Fernando R. Fernandez, Carmen C. Canavier, John. A. White

## Abstract

Gamma oscillations (40–140 Hz) play a fundamental role in neural coordination, facilitating communication and cognitive functions in the medial entorhinal cortex (mEC). While previous studies suggest that pyramidal-interneuron network gamma (PING) and interneuron network gamma (ING) mechanisms contribute to these oscillations, the precise role of inhibitory circuits remains unclear. Using optogenetic stimulation and whole-cell electrophysiology in acute mouse brain slices, we examined synaptic input and spike timing in neurons across layer II/III mEC. We found that fast-spiking interneurons exhibited robust gamma-frequency firing, while excitatory neurons engaged in gamma cycle skipping. Stellate and pyramidal cells received minimal recurrent excitation, whereas fast-spiking interneurons received strong excitatory input. Both excitatory neurons and fast-spiking interneurons received gamma frequency inhibition, emphasizing the role of recurrent inhibition in gamma rhythm generation. Notably, gamma activity was reduced, but persisted after AMPA/kainate receptor blockade, indicating that interneurons can sustain gamma oscillations independently through an ING mechanism. Selective activation of PV+ interneurons confirmed their ability to sustain fast gamma inhibition autonomously. To further assess the interplay of excitation and inhibition, we developed computational network models constrained by our experimental data. Simulations revealed that weak excitatory input to interneurons supports fast ING-dominated rhythms (∼100–140 Hz), while strengthening excitatory drive induces a transition to slower PING-dominated oscillations (60–100 Hz). These findings highlight the dominant role of inhibitory circuits in sustaining gamma rhythms, demonstrate how excitation strength tunes the oscillatory regime, and refine models of entorhinal gamma oscillations critical for spatial memory processing.

**Significance Statement:** Gamma oscillations in the medial entorhinal cortex (mEC) are essential for spatial navigation and memory, yet the mechanisms underlying their generation remain unresolved. Combining optogenetics, whole-cell electrophysiology, and computational modeling, we show that fast-spiking interneurons can autonomously sustain gamma rhythms via interneuron network gamma (ING). Blocking excitatory input reduced, but did not abolish gamma-frequency inhibition, and selective activation of PV+ interneurons confirmed their capacity to generate fast gamma independently. Modeling revealed that excitatory strength tunes the oscillatory regime, with weak excitation favoring fast ING and stronger excitation inducing slower pyramidal-interneuron network gamma (PING). These findings refine entorhinal gamma models and suggest a hybrid mechanism for switching between faster and slower gamma critical for spatial computation.

## Introduction

Cortical network oscillations play a fundamental role in coordinating neural activity (Buzsáki and Draguhn, 2004; Fries, 2009), facilitating communication between brain regions (Colgin, 2013), and supporting cognitive functions such as spatial navigation and memory (Buzsáki and Moser, 2013). In the medial entorhinal cortex (mEC), grid cells—neurons that exhibit spatially periodic firing—are strongly modulated by theta (4–12 Hz) and gamma (40–140 Hz) oscillations. Theta rhythms regulate the timing of grid cell firing (Fyhn et al., 2004; Hafting et al., 2005, 2008) and enable hippocampal-entorhinal interactions (Buzsáki and Moser, 2013). Gamma rhythms have been implicated in sensory processing, attention, and working memory (Colgin et al., 2009).

Gamma oscillations are hypothesized to arise from local circuit interactions involving both excitatory principal neurons and inhibitory interneurons. Two primary mechanisms have been proposed: pyramidal-interneuron network gamma (PING), which relies on excitatory-inhibitory interactions (Traub et al., 1996; Börgers and Kopell, 2003; Wang, 2010; Buzsáki and Wang, 2012), and interneuron network gamma (ING), which emerges from mutual inhibition among fast-spiking interneurons (Whittington et al., 2000; Bartos et al., 2007; Buzsáki and Wang, 2012). These models are not mutually exclusive and may operate in parallel or under different conditions. Moreover, there are different variants of PING. In classic PING, interneurons do not fire unless they receive phasic excitation (Börgers and Kopell, 2005; Tiesinga and Sejnowski, 2009; Börgers and Walker, 2013). Alternatively, interneurons can receive suprathreshold drive as in optogenetic stimulation of Thy1 expressing neurons (Pastoll et al., 2013), where they fire repetitively without input from the excitatory cells, therefore a different mechanism is required. Here, we distinguish between classic “driven inhibitory (I) cell” PING mechanism and our proposed “excitatory (E) cell recovers first” PING mechanism.

From previous models, E-I connectivity is sufficient in principle to account for theta-nested gamma oscillations and continuous attractor dynamics in EC (Pastoll et al., 2013; Solanka et al., 2015). The addition of I-I connections in models of EC can stabilize grid cell dynamics and increase the frequency of gamma oscillations (Solanka et al., 2015), suggesting that hybrid E-I-I networks can generate activity across the wide range of gamma frequencies observed in the mEC (Colgin et al., 2009).

Pharmacological evidence supports both PING and ING mechanisms. Blockade of AMPA/kainate receptors significantly reduces gamma power in the inhibitory currents of mEC stellate cells (Pastoll et al., 2013) and throughout the hippocampal–entorhinal system (Butler et al., 2018), consistent with PING. However, in the mEC, gamma oscillations appear to persist at reduced strength following AMPA/kainate receptor blockade, suggesting that inhibitory synchronization (i.e., ING) may also be sufficient.

Principal neurons in layer II/III of mEC include stellate and pyramidal cells (Alonso and Klink, 1993; Canto and Witter, 2012) with distinct electrophysiological profiles (Fig. 1-1). Grid cells, predominantly found among stellate and pyramidal neurons (Sargolini et al., 2006; Domnisoru et al., 2013), have been proposed to rely on recurrent excitatory connectivity (Fuhs and Touretzky, 2006; McNaughton et al., 2006). However, synaptic connectivity studies indicate that excitatory connectivity is sparse (Dhillon and Jones, 2000; Couey et al., 2013; Pastoll et al., 2013; Fuchs et al., 2016), particularly among stellate cells, but see (Winterer et al., 2017), raising the possibility that inhibitory circuits play a central role in shaping grid cell dynamics.

**Figure 1:**
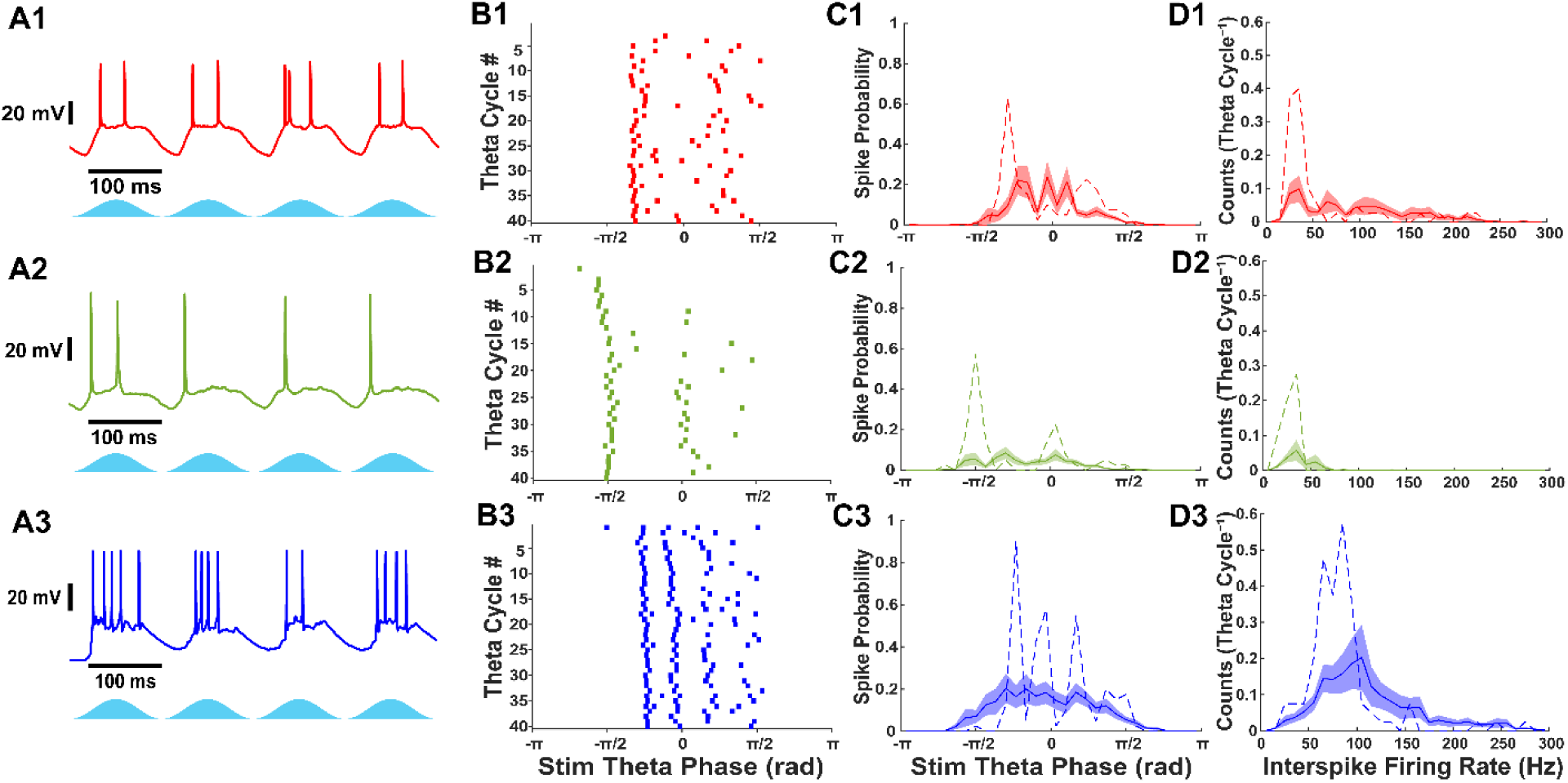
Stellate, pyramidal, and fast-spiking interneurons exhibit different firing patterns during theta frequency optogenetic stimulation of Thy1+ neurons. A) Current clamp recordings of (1) stellate cells, (2) pyramidal cells, and (3) fast-spiking interneurons during theta frequency optogenetic stimulation of Thy1+ neurons. B) Raster plots during 40 theta cycles of optogenetic stimulation from the cells shown in A. C) Average histogram of theta stimulation spike phase for stellate, pyramidal and fast-spiking interneurons. Examples from A and B are shown as dashed line. Shaded region indicates S.E.M. D) Average histogram of interspike firing rate distribution for stellate, pyramidal, and fast-spiking interneurons. Examples from A and B are shown as dashed line. Shaded region indicates S.E.M.

Parvalbumin-positive (PV+) fast-spiking interneurons, which constitute approximately half the inhibitory population in mEC (Wouterlood et al., 1995; Miettinen et al., 1996), also have a distinct electrophysiological profile (Fig. 1-1) and are critical for gamma rhythm generation (Sohal et al., 2009). These interneurons provide fast, perisomatic inhibition to grid cells (Buetfering et al., 2014; Miao et al., 2017), but have not been studied in detail (Sutton et al., 2024) One study reported strong chemical and electrical coupling (Fernandez et al., 2022) at distances < 150 μm, whereas another found only gap junctions at distances > 100 μm (Huang et al., 2024). Clarifying I-I connectivity is essential to understanding whether PV+ interneurons can generate gamma oscillations without gamma-frequency excitation.

In this study, we use optogenetic stimulation, whole-cell recordings, pharmacological manipulations, and computational modeling to study the contributions of fast-spiking interneurons in the mEC to gamma oscillations. We compare the timing and strength of inhibitory input across cell types, assess interspike intervals and phase-locking to theta-frequency drive, and evaluate the persistence of gamma oscillations following blockade of AMPA/kainate receptors. In parallel, we develop network simulations to examine how excitatory-inhibitory balance regulates gamma rhythms under conditions of suprathreshold drive to PV+ interneurons. These combined approaches allow us to study how excitation strength tunes the transition between underlying mechanisms.

## Materials and methods

All procedures were approved by the Institutional Animal Care and Use Committee of Boston University and conformed to NIH guidelines. Adult (2–6 months old) male and female mice were used in roughly equal numbers. A transgenic mouse line constitutively expressed ChR2 in both excitatory and inhibitory cells under the Thy1 promoter (Thy1-ChR2-EYFP, JAX strain #007612). A separate transgenic mouse line expressed Cre recombinase under the PV promoter (PV-Cre, JAX strain #017320). Parvalbumin is expressed almost exclusively in fast-spiking inhibitory interneurons in mEC. The transgenic PV-Cre mouse line was cross bred with transgenic LoxP-ChR2-EYFP mice (JAX, strain #024109) to express ChR2 in PV+ fast-spiking interneurons (PV-ChR2) after Cre-Lox recombination.

### Acute slice preparation

All measurements were obtained from 400-micron thick horizontal mouse brain slices. Bilateral slices from the dorsomedial mEC (∼3.2–4.3 mm from the dorsal surface of the brain) were used. Mouse brains were extracted after isofluorane overdose and decapitation. During slicing, the brain was submerged in sucrose-substituted artificial cerebrospinal fluid (ACSF) solution (in mM: sucrose 225, KCl 2.5, NaH_2_PO_4_ 1.25, MgCl_2_ 3, NaHCO_3_ 25, glucose 20, and CaCl_2_ 0.5) at 4°C that was continuously perfused with 95% oxygen / 5% carbon dioxide (carboxy) gas and sliced with a vibratome (VT1200, Leica Microsystems). Then, brain slices are moved to a separate chamber containing standard ACSF (in mM: NaCl 125, NaHCO_3_ 25, D-glucose 25, KCl 2.5, CaCl_2_ 2, NaH_2_PO_4_ 1.25, and MgCl_2_ 1) perfused with carboxy gas and incubated at 37°C for 30 minutes. After incubation, the brain slices are allowed to recover for 15 minutes at room temperature (∼20°C) where they remained until being used for whole-cell electrophysiology.

### Whole-cell electrophysiology

For electrophysiology experiments, brain slices were continuously perfused with ACSF at a temperature of 35–37°C. The ACSF was bubbled with 95%/5% carboxy gas throughout the experiments. Borosilicate glass pipettes were pulled (Sutter Instrument P-97) and filled with intracellular fluid solution (in mM: K-gluconate 136, KCl 4, HEPES 10, diTrisPhCr 7, Na_2_ATP 4, MgCl_2_ 2, TrisGTP 0.3, and EGTA 0.2, and buffered to pH 7.3 with KOH) and pipettes with resistances of 4–8 MΩ were used for electrophysiology recordings. Pipette offset was compensated prior to achieving on-cell patch recordings. Once the patch electrode was sealed to the cell membrane (>1 GΩ seal), pipette capacitance was compensated. Series resistances < 40 MΩ were used in this study with changes < 20% throughout recordings. For voltage clamp recordings, series resistance was compensated 50-70%. Using voltage clamp, excitatory and inhibitory post-synaptic currents were recorded at −70 and 0 mV, respectively. For our recording conditions, the reversal potential for chloride is −75 mV. For current clamp recordings, full bridge balance compensation was used. Liquid junction potentials were not corrected. Electrophysiology data was amplified using Axon Instruments MultiClamp 700B and sampled at 30 kHz using Axon Instruments DigiData 1440A. Custom protocols were designed using pClamp 7.0 software to control data collection and optogenetic stimulation. Pipettes and cells were visualized with diffuse interference contrast.

### Optogenetic stimulation

A 470 nm LED (Thorlabs, M470L4) delivered widefield optogenetic stimulation through a 40x objective lens with a typical intensity of ∼2 mW/mm^2^, although up to 24 mW/mm^2^ was used to probe connectivity. For each cell, the same light power was used throughout recordings (i.e., voltage clamp at −70 mV and 0 mV; current clamp at 0 pA). However, the light power was adjusted for each cell (i.e., field-of-view) to drive sufficient activity for gamma frequency synchronization in the surrounding local network. The optimal light power varied to correct possible variations in slice excitability and functional connectivity. The light intensities used in our study across all recordings have no linear relationship with the gamma power and frequency of inhibition, or firing rate of mEC neurons in Thy1 experiments (Fig. 3-1). During PV stimulation, we specifically tested the relationship between light intensity and gamma frequency/power among individual neurons using a 100 ms pulse as opposed to a sinusoidal waveform (Fig. 3-1). We found that the gamma frequency and power of PV inhibition increase over low light intensities, then plateaus as light intensity approaches the median intensities used for each cell type.

### Pharmacology

During pharmacological experiments, the AMPA/kainate (5 µM DNQX disodium, Tocris Bioscience Cat. No. 2312) or GABA_A_ (10 µM SR995531 hydrobromide (Gabazine), EMD Millipore Corp.) receptor antagonist was bath-applied, and recordings were repeated after a confirmed wash-in period sufficient to fully block either AMPA/kainate or GABA_A_ currents. To determine the required duration, spontaneous EPSCs or IPSCs were monitored in fast-spiking interneurons during DNQX or Gabazine wash-in, respectively, allowing us to observe the suppression and eventual elimination of AMPA/kainate or GABA_A_ mediated events in real time. Complete blockade was consistently achieved by ∼10 minutes, and this standardized wash-in period was subsequently used for all recordings.

### Cell type classification

The major electrophysiological cell types in the mEC were statistically separated by their electrophysiological properties (Fig. 1-1). Current steps were injected from −200 pA to 525 pA at 25 pA intervals to characterize the subthreshold and firing properties of each cell. Putative stellate and pyramidal cells in layer II/III mEC were classified based on membrane sag potential and membrane time constant as used previously (Fernandez et al., 2022). Fast-spiking interneurons were easily identified by their discontinuous membrane current – spike rate relationship, high threshold firing, burst firing, fast membrane time constant, and short spike half width.

### Data analysis and statistics

All data analysis was performed with custom written algorithms in MATLAB. Action potential peaks were detected and registered to the phase of the stimulation period. Spike phase histograms were computed with 30 equally spaced bins over the stimulation period (width = π/15 radians) and were normalized by the number of theta cycles (40 cycles for each cell). Interspike frequency histograms were computed with 10 Hz bins and the bin counts were normalized by the number of theta cycles (40 cycles for each cell).

For current recordings, raw data was forward and reversed filtered from 50–200 Hz with 4th order butterworth filters. To analyze the frequency and amplitude of gamma oscillations, the continuous wavelet transform was computed in MATLAB. The analytic morlet wavelet (ω_0_ = 6 rad/sec) was used to calculate the scalogram for each theta cycle with 32 scales per octave. The network activity during the first stimulation period varied greatly compared to subsequent cycles and was removed from analysis. The magnitude of the scalogram was averaged over the next 40 theta cycles to determine the peak gamma frequency, phase and power of the membrane currents for each cell. Any theta cycles with large current artifacts or spikes (> 3000 pA) were removed from analysis. Low average peak gamma power recordings (< 20 pA^2^ for Thy1 and < 10 pA^2^ for PV experiments), scalograms peaks with large bandwidths (> 100 Hz for Thy1 and > 110 Hz for PV experiments), and scalograms with peak powers less than five multiples above the average (SNR < 5) were removed for accurate gamma frequency comparisons. Total gamma power was summed over the 60-140 Hz range for all theta phases in the average scalogram within the cone of influence to remove edge effects in wavelet analysis.

Non-parametric methods were used for statistical testing to control for non-normality and differences in sample sizes unless otherwise noted. For paired recordings obtained before and after the application of DNQX, the two-sided Wilcoxon signed-rank test was used to compare the change in peak gamma power or frequency, whereas the paired t-test was used to compare changes in total gamma power. The Kruskal-Wallis test was used for comparisons between independent cell types. A significant effect was followed by a post-hoc Dunn’s test with Bonferroni correction for multiple comparisons. Significance was determined at p < 0.05. Data was reported as mean ± S.E.M. for parametric statistical comparisons or median (Q1 – Q3) for non-parametric unless otherwise noted.

### Computational methods

All simulations were carried out using the NetPyNE (Dura-Bernal et al., 2019) simulator based on Python. The network consists of 500 single compartment model neurons of two populations: 100 inhibitory PV+ interneurons and 400 excitatory stellate cells. The optogenetic theta drive was simulated by a sinusoidal 8 Hz conductance waveform biased to have only positive values with a peak of 4 nS for the PV cells and 3 nS for the stellate cells and a reversal potential of 0 mV. Five readout neurons from each population were voltage-clamped at 0 mV to measure inhibitory currents during theta drive. The frequency and power of oscillations in the inhibitory currents were analyzed using the analytic morlet wavelet in the neuro digital signal processing package of Python to obtain the scalogram.

Since a previous model (Pastoll et al., 2013; Solanka et al., 2015) also addressed optogenetically driven theta, we point out some essential differences in our models. Differences include: 1) the optogenetic drive in their model is a current whereas in ours it is a conductance that reverses at 0 mV, 2) our interneurons are biophysically calibrated to capture the full breadth of heterogeneity in their active and passive properties, 3) our simulations include both chemical and electrical interconnectivity between interneurons, and 4) the optogenetic drive is subthreshold in their model and suprathreshold in ours.

*PV+ interneuron model*: The model consisting of 100 PV+ interneurons is based on our previous work (Via et al., 2022), calibrated using data based on the passive and intrinsic properties of these neurons (Fernandez et al., 2022).

The single compartment model has five state variables: the membrane potential, *V*, and four gating variables (m, h, n, and a) which use the same kinetic equations as the original Hodgkin-Huxley model (Hodgkin and Huxley, 1952), but with different parameters tuned to replicate the dynamics of fast-spiking interneurons in the medial entorhinal cortex and given in Table 1 of (Via et al., 2022). We included two delayed rectifier K^+^ currents (I_Kv1_ and I_Kv3_). The differential equation for the membrane potential V is given by Eq. 1:

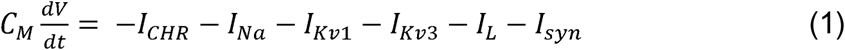

where C_M_ is the membrane capacitance, I_CHR_ is the simulated sinusoidal current replicating the optogenetic 8 Hz (theta) drive normally distributed with a mean-conductance of 4 nS and standard deviation of 0.4 nS, I_Na_ is the fast sodium current, I_L_ is the leak current, I_syn_ is the GABA_A_ synaptic current. The ionic-current equations (Eq. 2-5) are:

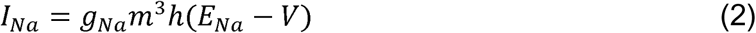

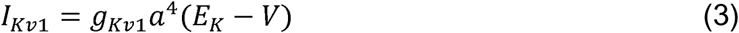

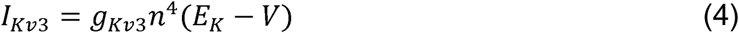

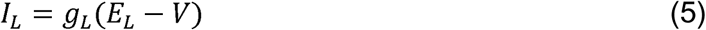

with E_Na_ = 50 mV, E_K_ = −90 mV and E_L_ = −65 mV. The dynamics of the gating variables are given by 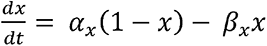 for the activation variables (m, n, a) and by 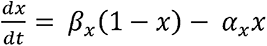 for the inactivation variable h, where *α_x_* = *k*_1*x*_ (*θ_x_* −*V*)/(exp((*θ_x_* −*V*)/*σ*_1*x*_) −1) and *β_x_* = *k*_2*x*_ exp(*V* /*σ*_2*x*_) with parameters in Table 1.

**Table 1:**
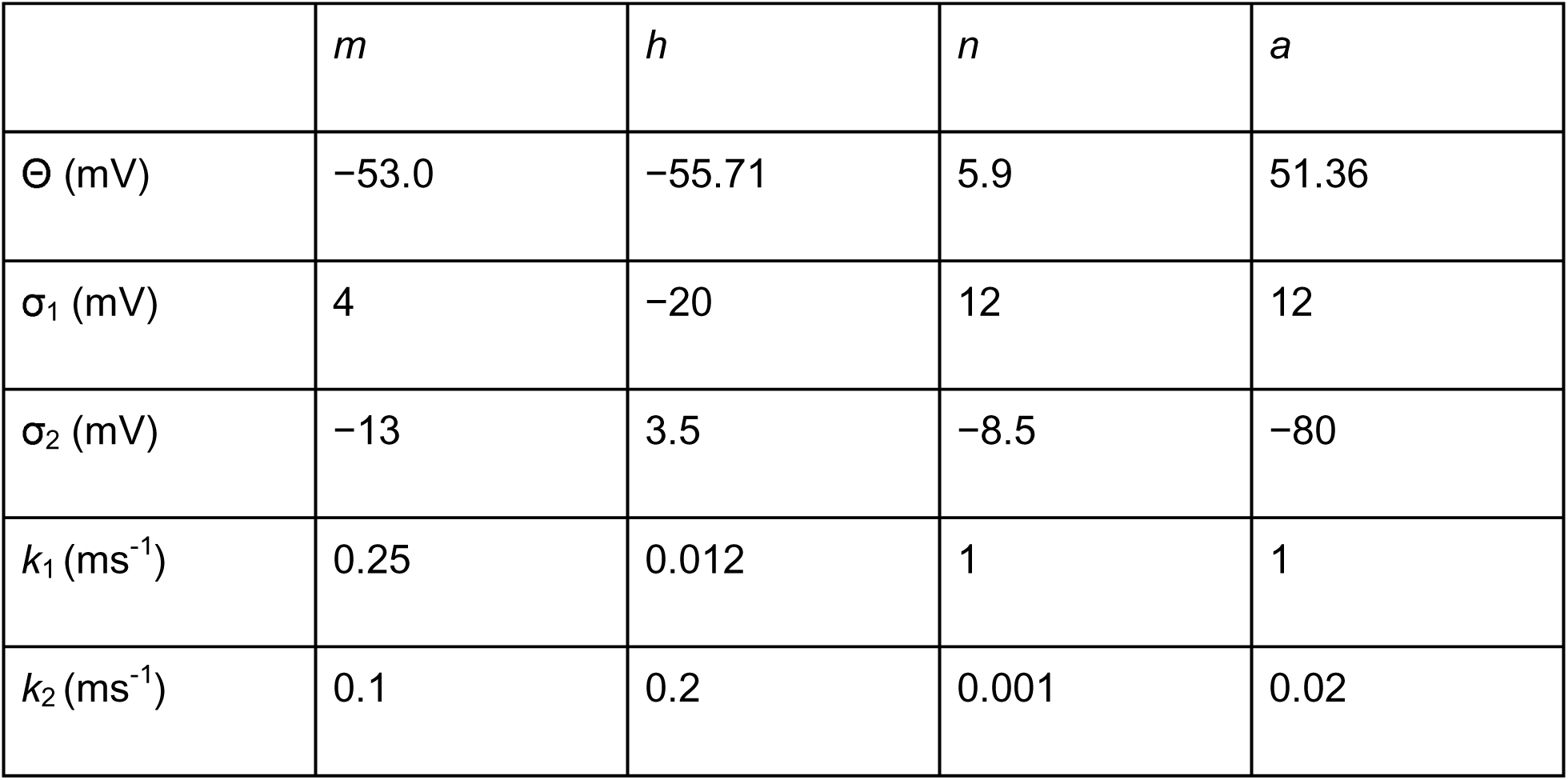
Parameters for gating variables in Via et al. (2022) model.

| | $m$ | $h$ | $n$ | $a$ |
| --- | --- | --- | --- | --- |
| $\Theta$ (mV) | -53.0 | -55.71 | 5.9 | 51.36 |
| $\sigma_1$ (mV) | 4 | -20 | 12 | 12 |
| $\sigma_2$ (mV) | -13 | 3.5 | -8.5 | -80 |
| $k_1$ (ms<sup>-1</sup>) | 0.25 | 0.012 | 1 | 1 |
| $k_2$ (ms<sup>-1</sup>) | 0.1 | 0.2 | 0.001 | 0.02 |

*E cell model*: The E cell model ) is based on the original Hodgkin-Huxley conductance-based model modified to better capture the firing properties of the stellate cells in the mEC (Alonso and Klink, 1993). Heterogeneity is introduced by jittering the conductance values from a Gaussian distribution to obtain 400 stellate cell models, and by introducing conductance noise to these cells based on the Ornstein-Uhlenbeck (OU) stochastic process (Fellous et al., 2003). One process was used for the excitatory noise (E-Noise) and one for inhibitory noise (I-Noise) with the reversal potentials 0 mV and −75 mV respectively. We used 0.9 nS as mean conductances for both and 0.02 nS as standard deviation. Time constant values were 2.728 ms and 10.49 ms for the I-Noise and E-Noise respectively.

*Connectivity*: The probability of I->I connections was 30%. The probability of I->E was 40% and E->I connectivity was 30%. The weights of the I->E connections were also randomized using a lognormal distribution as in (Fernandez et al., 2022) scaling the mean by a factor of 2.5. We used a probability of 18% for each pair of PV-IN to have gap junctional connectivity. To ensure that the results were reproducible, and not an artefact of the connectivity parameters for a single simulation, the simulations varying the E->I conductance strength were run with 15 different seeds to randomize the connectivity. The simulations varying E->I and I->E connections simultaneously, were run with 9 different seeds that randomize connectivity.

*Synapses*: As in the Via model (Via et al., 2022), we modeled both electrical and chemical synapses for the I->I connections. We calibrated the inhibitory synapses onto each other with a rise time constant of 0.3 ms, a decay time constant of 4 ms. For most of the analysis (except for Fig. 7B), we used the synaptic reversal potential in I->I connections as E_GABA_ = −75 mV for the chemical synapses. Inhibitory synapses onto the E cells were calibrated with a rise time constant of 0.4 ms and decay time constant of 6 ms, with E_GABA_ = −65 mV (Sauer et al., 2012). The E cells in this model are not connected to each other, based on the experimental data (Dhillon and Jones, 2000; Couey et al., 2013; Pastoll et al., 2013; Fuchs et al., 2016). Excitatory synapses to the PV+ interneurons use E_AMPA_ = 0 mV and an exponential synapse with a decay time constant of 1 ms. All the chemical synaptic connections were modeled with randomized synaptic delay following a uniform distribution of 0.6-1 ms.

*Code Accessibility:* The network model and associated programs to run simulations is freely available online at https://github.com/AnanthVS23/mEC_Network_Simulations. The code is available as Extended Data. The simulations and models in this paper were run on the LSUHSC-NO Tigerfish HPC Cluster, using NSF Grant #2018936. Tigerfish runs the Linux CentOS operating system and has a total of 1440 compute cores providing over 193 teraFLOPS of computing power. The current cluster configuration has 36 computing nodes each with 40 compute cores and 192 GB of RAM. There is also a large memory node with access to 1500 GB of RAM and a GPU node with access to 192 GB of RAM. Secondary storage for the cluster consists of 576 TB of usable storage in a BeeGFS file system along with a 45.6 TB SSD Layer to accelerate performance. The system was built by Advanced Clustering Technologies and is expandable to accommodate future upgrades if the need arises and funding is available. We have not used GPUs for our simulations.

## Results

A transgenic mouse line was used to express ChR2 primarily in stellate cells and fast-spiking interneurons in layer II/III mEC. However, pyramidal neurons have also been shown to express ChR2 in mEC with the Thy1 promoter (Proskurina and Zaitsev, 2021). Acute brain slices were prepared from roughly equal numbers of adult male and female Thy1-ChR2-EYFP mice. Whole-cell patch clamp recordings were obtained from stellate, pyramidal and fast-spiking interneurons in layer II/III mEC during 8 Hz theta-frequency sinusoidal optogenetic stimulation, to simulate network drive observed in spatial navigation. A previous study investigated the responses of stellate cells and fast-spiking interneurons during theta frequency drive in Thy1-ChR2 mice (Pastoll et al., 2013). In Figures 1-4, we aim to replicate the prior findings and provide additional measurements in pyramidal cells. We also provide separate functional connectivity measures of excitation and inhibition in all cell types that are essential for building accurate models of the mEC and spatial processing.

**Figure 2:**
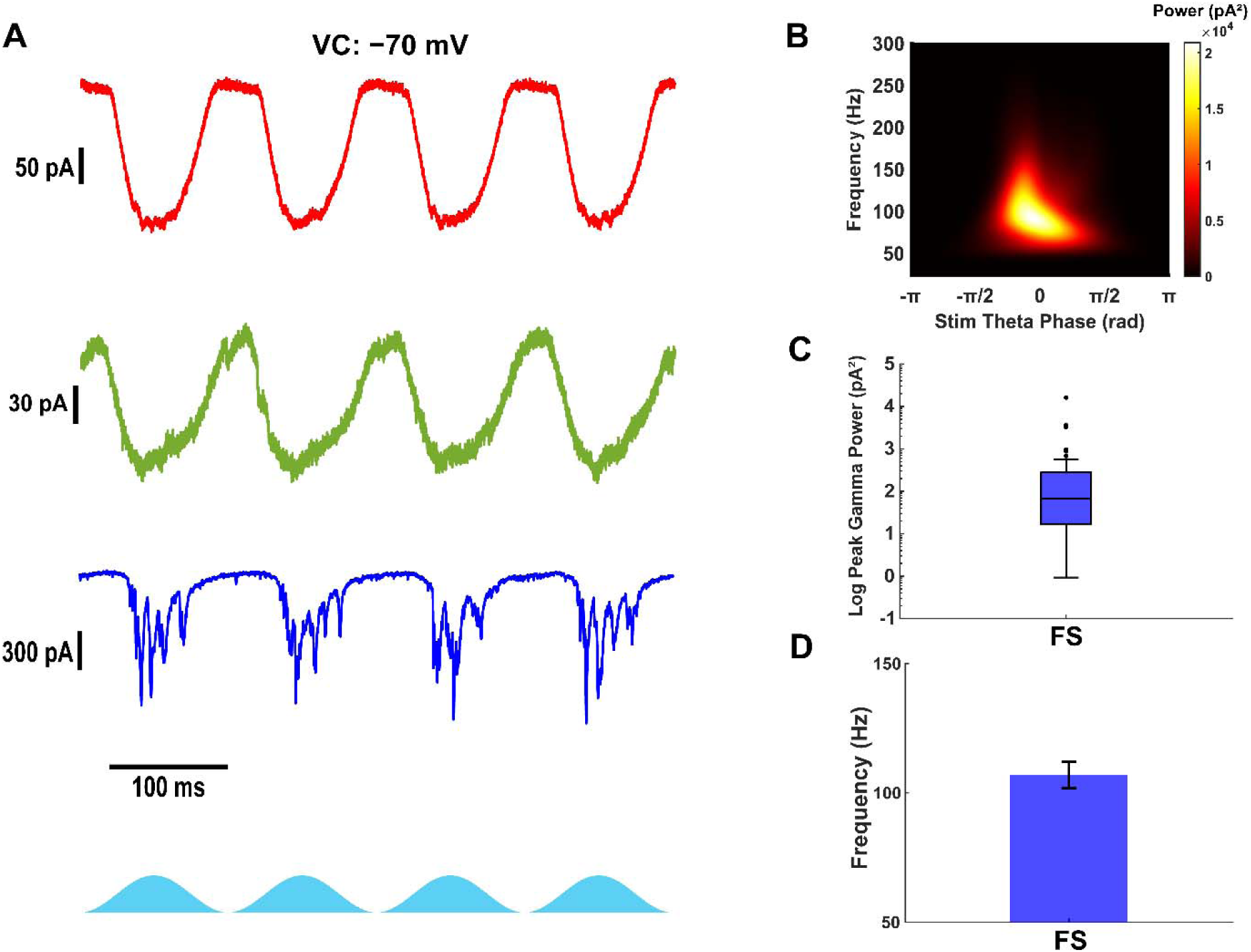
Only fast-spiking interneurons receive fast and strong gamma frequency excitatory currents. A) Example excitatory currents from stellate (top, red), pyramidal (middle, green) and fast-spiking interneurons (bottom, blue) during theta-frequency optogenetic stimulation of Thy1+ neurons. B) Average scalogram of gamma frequency excitatory currents in fast-spiking interneuron from A during 40 theta cycles of optogenetic stimulation. C) Peak gamma power from scalograms of excitatory currents in fast-spiking interneurons. Filled circles indicate outliers. D) Gamma frequency at the peak gamma power from scalograms of excitatory currents in fast-spiking interneurons.

**Figure 3:**
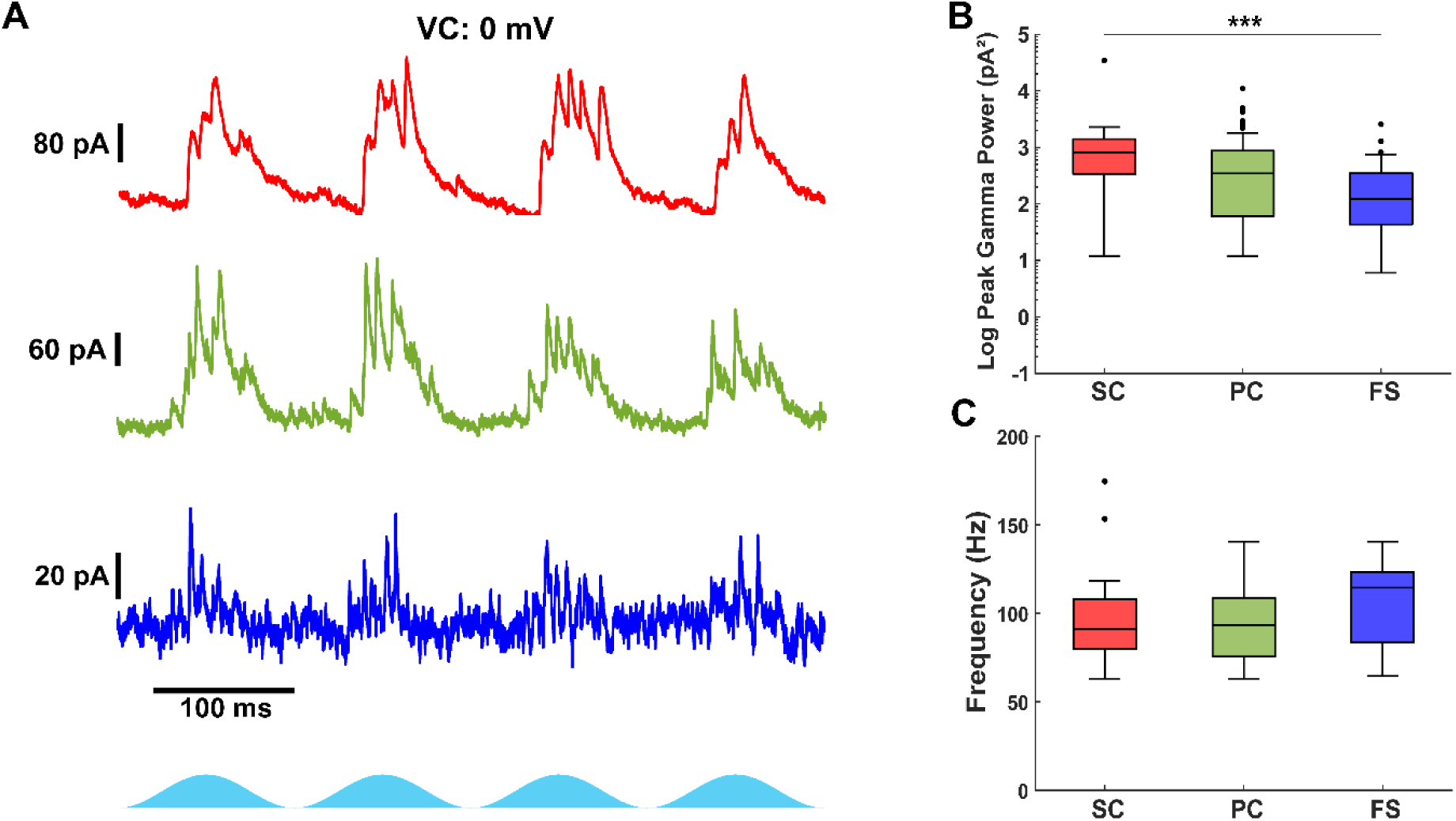
Excitatory neurons receive stronger gamma frequency inhibitory input than fast-spiking interneurons. A) Example inhibitory current recordings in stellate (top, red), pyramidal (middle, green) and fast-spiking interneurons (bottom, blue) during optogenetic stimulation of Thy1+ neurons. B) Comparison of the peak gamma power from the average scalogram of inhibitory currents received by each cell. Gamma frequency inhibition is weaker in fast-spiking interneurons. C) Comparison of the peak gamma frequency from the average scalogram of excitatory currents received by each cell. The gamma frequency of inhibitory currents are similar across cell types. Filled circles indicate outliers.

**Figure 4:**
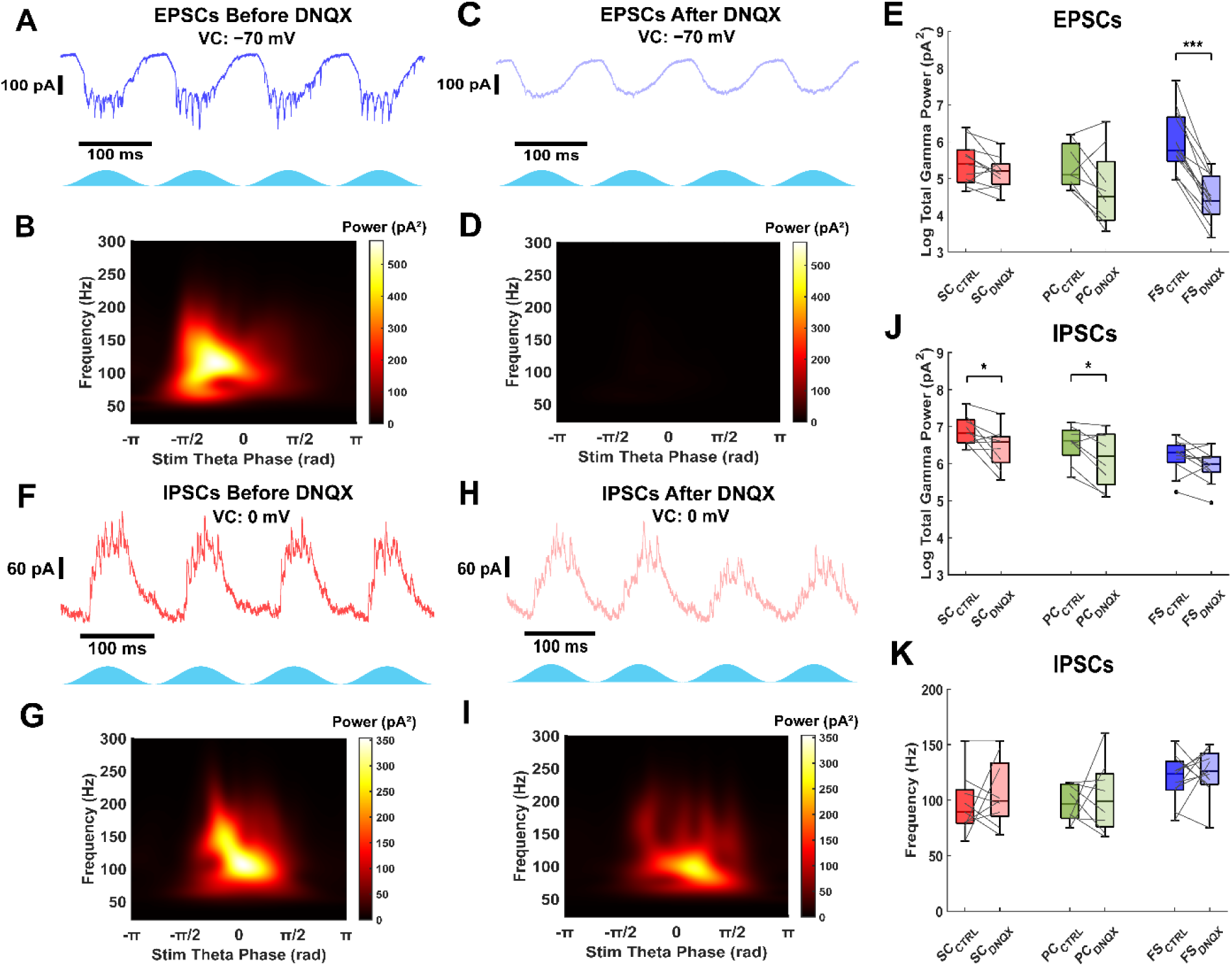
Thy1+ interneurons provide fast and strong inhibition in the absence of fast excitatory synaptic input. A) Example excitatory current trace in fast-spiking interneuron during Thy1 stimulation. B) Average scalogram from 40 theta cycles of excitatory currents from example trace in A. Data was filtered from 50 to 200 Hz. C) Excitatory currents recorded from the same fast-spiking interneuron in A after blocking AMPA-mediated synaptic input with 5 µM DNQX. D) Average scalogram from 40 theta cycles of excitatory currents from example trace in C. Data was filtered from 50 to 200 Hz. E) Comparison of excitatory current peak gamma power for all cell types before and after DNQX. Gamma frequency currents are abolished in fast-spiking interneurons. F) Example inhibitory current trace in stellate cell before blocking AMPA receptors with DNQX. G) Average scalogram from 40 theta cycles of inhibitory currents from example trace in F. H) Inhibitory currents recorded from the same stellate cell in F after blocking AMPA receptors with DNQX. I) Average scalogram from 40 theta cycles of inhibitory currents from trace in H. J) Comparison of total inhibitory current gamma power for all cell types before and after DNQX. The total inhibitory current gamma power is reduced but not abolished. K) Comparison of inhibitory current gamma frequency for all cell types before and after DNQX. The frequency of inhibition doesn’t significantly change with synaptic excitation blocked. Gray lines indicate paired data points.

### Firing characteristics of stellate, pyramidal and fast-spiking interneurons during theta-frequency Thy1+ stimulation

Voltage recordings were analyzed to characterize the firing responses of different neuron types during optogenetic stimulation of Thy1+ neurons (Fig. 1A). All cell types exhibited firing activity during stimulation. However, pyramidal neurons demonstrated a lower probability of activation (77.1%, n = 27/35) compared to stellate (94.4%, n = 17/18) and fast-spiking interneurons (89.4%, n = 17/19). The firing rates varied among the neuron types. Stellate and pyramidal neurons fired an average of 1.54 ± 0.28 and 0.67 ± 0.12 spikes per theta cycle, respectively, while fast-spiking interneurons exhibited a higher firing rate of 2.30 ± 0.56 spikes per theta cycle. While most fast-spiking interneurons fired at least once during optogenetic Thy1 stimulation, the consistency of the firing responses were quite variable (Spikes per theta cycle median: 1.10, Q1: 0.44, Q3: 4.08, min: 0, max: 7.28, n = 19), indicating that not all fast-spiking interneurons were strongly driven by the optogenetic stimulus in Thy1-ChR2 mice. Previously, Pastoll et al. (2013) measured the firing rates of stellate cells and fast-spiking interneurons during theta frequency stimulation in layer II of Thy1-ChR2 mice. The firing rates of stellate cells in our study are similar to those measured in Pastoll et al. (2013), and the resting membrane potentials of stellate cells (−63 ± 3 mV, mean ± s.d., n = 24 cells from 15 mice) appear similar to those reported by the same lab (Pastoll et al., 2020). Fast-spiking interneurons fired more often in their study (Pastoll et al., 2013), however, the average firing rates of mEC neurons in our study are within the range of in vivo measurements in layer II mEC (Chrobak and Buzsáki, 1998). In addition, we found that Thy1 stimulation activated most pyramidal cells, consistent with a histological study that found expression of ChR2 in mEC pyramidal cells of Thy1-ChR2 mice (Proskurina and Zaitsev, 2021), whereas Pastoll et al. (2013) reported no activation in this cell type.

The spike phase distribution for stellate cells showed peaks corresponding to fast gamma frequencies (Fig. 1C), and individual stellate cells occasionally fired at ∼100 Hz (Fig. 1D). However, most stellate neurons displayed interspike intervals that were slower multiples of the dominant fast gamma frequency, indicative of gamma cycle skipping. Pyramidal neurons, in contrast, were unable to fire at fast gamma frequencies and instead skipped one or two gamma cycles between action potentials. Fast-spiking interneurons exhibited a broader spike-phase distribution, with some firing multiple times per putative gamma cycle, while the majority fired at fast gamma frequencies. These results highlight distinct cell-type-specific firing dynamics in response to theta-frequency stimulation, with fast-spiking interneurons most consistently engaging in fast gamma activity, stellate cells exhibiting intermittent participation and cycle skipping, and pyramidal neurons showing limited involvement, suggesting differential contributions to gamma rhythm generation in layer II/III mEC.

### Fast-spiking interneurons receive strong gamma frequency excitatory input during Thy1+ stimulation

Previous studies have suggested that stellate cells have minimal recurrent connectivity in layer II mEC (Dhillon and Jones, 2000; Couey et al., 2013; Pastoll et al., 2013), while pyramidal neurons exhibit slightly higher connectivity (Fuchs et al., 2016). Notably, fast-spiking interneurons receive gamma-frequency excitatory currents during theta-frequency optogenetic stimulation of Thy1+ neurons (Pastoll et al., 2013). However, these measurements included a mix of excitatory and inhibitory currents, as voltage was clamped at −50 mV. To isolate fast excitatory inputs, we performed voltage-clamp recordings at −70 mV, thereby minimizing inhibitory and excluding NMDA-mediated currents while capturing AMPA-mediated and ChR2 currents.

Our recordings confirmed that fast-spiking interneurons received AMPA-mediated excitatory currents at fast gamma frequencies in addition to theta-frequency ChR2 currents (Fig. 2A, 2B) in accordance with (Pastoll et al., 2013). Whereas all stellate cells exhibited ChR2-induced excitatory currents, they did not display fast inward currents during Thy1+ stimulation, indicating negligible recurrent excitation. Due to strong inhibitory input and imperfect space clamp, outward currents were partially observed even at −70 mV. Most pyramidal cells expressed ChR2 currents and small excitatory post-synaptic currents (EPSCs), but these synaptic events lacked robust oscillations. In contrast, many fast-spiking interneurons received strong gamma-frequency excitation during optogenetic stimulation (Peak gamma power: 68 pA^2^ (17 – 281), n = 38, Fig. 2C; Gamma frequency: 106.9 ± 5.1 Hz, n = 19/38, Fig. 2D), suggesting that they are uniquely recruited by local excitatory cells and well-positioned to provide gamma frequency inhibition, as demonstrated by their firing rates in Fig. 1. Recordings with low gamma power, low signal-to-noise, and high spectral bandwidth were removed from frequency analysis (see Materials and methods).

### Excitatory cells receive stronger inhibition than fast-spiking interneurons during Thy1+ stimulation

Fast-spiking interneurons are key generators of gamma frequency activity, but how different excitatory and inhibitory cell types receive and integrate inhibitory input remains unclear. Given the distinct connectivity patterns and intrinsic properties of stellate, pyramidal, and fast-spiking interneurons, we sought to determine how inhibitory input varies across these cell types during optogenetically driven network activity.

To compare inhibitory input across neuron types, we voltage-clamped neurons at 0 mV to minimize excitatory synaptic and ChR2 currents. All cell types received fast gamma frequency inhibitory currents (Fig. 3A). Cell type had a significant effect on the peak inhibitory gamma power (χ^2^ = 14.22, p = 8.2e−4, Kruskal–Wallis test). Stellate cells received stronger inhibitory gamma currents than fast-spiking interneurons (Stellate: 823 pA^2^ (339 – 1399), n = 20; Fast-spiking: 124 pA^2^ (44 – 352), n = 34; p = 5.8e−4; Fig. 3B), but their inhibition was comparable to that of pyramidal neurons (Pyramidal: 346 pA^2^ (60 – 870), n = 74; p = 0.079). The inhibitory current power was nearly significantly different between pyramidal neurons and fast-spiking interneurons (p = 0.053). In general, stellate and pyramidal cells exhibited greater summation of inhibitory currents per theta period compared to fast-spiking interneurons. However, cell type did not have a significant effect on the frequency of inhibitory currents (χ²_(2,95)_ = 5.06, p = 0.079, Kruskal–Wallis test). Therefore, fast-spiking interneurons received inhibition at a similar frequency compared stellate (Fast-spiking: 114.4 Hz (83.7 – 123.4), n = 16/34; Stellate: 91.1 Hz (80.0 – 107.8), n = 19/20; Fig. 3C) and pyramidal neurons (Pyramidal: 93.1 Hz (75.4 – 108.4), n = 63/74). These findings confirm that stellate cells receive gamma frequency inhibition in accordance with Pastoll et al. (2013). Further, we demonstrate that pyramidal cells and fast-spiking interneurons also receive gamma frequency inhibition, although excitatory neurons integrate stronger and more prolonged inhibition compared to fast-spiking interneurons.

A high degree of variability was observed in the power and frequency of gamma inhibition received by excitatory cells. One possible source of variability in the inhibition could be due to differences in the topographic distribution of fast-spiking interneurons along the dorso-ventral axis of the mEC (Beed et al., 2013). To control possible differences across the dorsal-ventral axis, we recorded inhibitory currents in a subset of excitatory neurons from the most dorsal and ventral slices of the mEC. We found no statistical differences in the peak power of gamma inhibition (Dorsal log power: 2.60 ± 0.22, n = 16; Ventral log power: 2.57 ± 0.19, n = 17; p = 0.90, t-test; Fig. 3-2A), peak gamma frequency (Dorsal frequency: 97.3 Hz (80.0 – 108.4), n = 15/16; Ventral frequency: 80.0 Hz (75.0 – 98.3), n = 17/17; p = 0.24, Wilcoxon rank sum test; Fig. 3-2B), or firing rate (Dorsal spikes per theta cycle: 0.93 (0.06 – 1.98), n = 12; Ventral spikes per theta cycle: 0.48 (0.05 – 0.78), n = 11; p = 0.31, Wilcoxon rank sum test; Fig. 3-2C) of excitatory cells across the dorsal and ventral extents of the mEC. While we found no statistical difference in the inhibition frequency due to similarly high variability in both the dorsal and ventral regions, there is a trend of higher frequency activity in the dorsal mEC. Future studies could investigate whether a possible gradient in the inhibitory frequency exists across the dorsal-ventral axis.

### Fast-spiking interneurons generate gamma frequency inhibition without AMPA-mediated excitation

Models of grid cells and theta-nested gamma oscillations in the medial entorhinal cortex sometimes assume a role for recurrent excitation. However, most previous connectivity studies have shown that stellate cells exhibit minimal recurrent excitation (Dhillon and Jones, 2000; Couey et al., 2013; Pastoll et al., 2013), while pyramidal cells display a higher probability of recurrent connectivity (Fuchs et al., 2016). As a result, alternative grid cell models propose that principal neurons are connected exclusively through recurrent inhibition. It is crucial to quantify the extent of AMPA-mediated excitation in layer II/III of the mEC to refine grid cell models.

To isolate AMPA-mediated excitation, we recorded excitatory currents before and after applying the AMPA receptor antagonist DNQX (5 µM). Blocking AMPA receptors abolished strong gamma-frequency excitatory currents in fast-spiking interneurons (Fig. 4A-E). Peak gamma power in excitatory currents significantly decreased in fast-spiking interneurons (Median log difference: −1.48 (−1.85 – −1.17), n = 13, p = 2.4e−4, Wilcoxon signed-rank test), while it remained unchanged in pyramidal neurons (Median log difference: −0.98 (−1.19 – −0.01), n = 8, p = 0.11, Wilcoxon signed-rank test) and stellate cells (Median log difference: −0.35 (−0.50 – 0.03), n = 11, p = 0.10, Wilcoxon signed-rank test). The cell type had a significant effect on the change in peak gamma power (χ²_(2,29)_ = 15.69, p = 3.9e−4, Kruskal–Wallis test). The reduction in peak gamma power for fast-spiking interneurons was significantly different compared to stellate (p = 4.0e−4) and pyramidal (p = 0.033) cells, but no significant difference was found between stellate and pyramidal cells (p = 1). These findings indicate that fast-spiking interneurons, but not stellate or pyramidal cells, receive local AMPA excitatory inputs, consistent with connectivity studies and Pastoll et al. (2013).

Pastoll et al. (2013) previously demonstrated that gamma frequency inhibitory currents in stellate cells significantly decreased after blocking both AMPA and NMDA excitatory currents but remained unchanged when only NMDA currents were blocked. This suggests that AMPA excitation plays a crucial role in sustaining strong gamma frequency activity in the mEC. To determine whether AMPA-mediated excitation is necessary for gamma frequency inhibition, we applied DNQX (5 µM) and recorded synaptic inhibitory currents before and after drug application.

During baseline theta-frequency optogenetic stimulation, all cell types received gamma-frequency inhibitory currents (e.g., Fig. 4F, G), as in our previous findings (Fig. 3). After blocking AMPA-mediated excitation, gamma-frequency inhibitory oscillations persisted across all cell types (Fig. 4H-K). Peak gamma power did not change significantly in stellate cells (Median log difference: −0.15 (−1.01 – 0.02), n = 9, p = 0.10, Wilcoxon signed-rank test), pyramidal neurons (Median log difference: −0.41 (−0.81 – −0.07), n = 8, p = 0.055, Wilcoxon signed-rank test), or fast-spiking interneurons (Median log difference: −0.23 (−0.51 – 0.09), n = 11, p = 0.21, Wilcoxon signed-rank test; data not shown). Although, total gamma power (60-140 Hz) analysis reveals a statistically significant decrease in the inhibitory currents received by stellate (Mean log difference: −0.44 ± 0.17, n = 9, p = 0.045, paired t-test) and pyramidal (Mean log difference: −0.39 ± 0.14, n = 8, p = 0.039, paired t-test) cells after AMPA receptor block, but not fast-spiking interneurons (Mean log difference: −0.23 ± 0.11, n = 11, p = 0.070, paired t-test; Fig. 4J), possibly explained by their comparatively weaker inhibition. The total gamma power reduction of inhibitory currents in stellate cells was previously observed (Pastoll et al., 2013). Additionally, the median frequency of these inhibitory oscillations remained unchanged across all groups (Median difference: Stellate: 0 Hz (−17.58 – 30.98), n = 9, p = 0.74; Pyramidal: −4.53 Hz (−21.99 – 28.21), n = 8, p = 0.95; Fast-spiking: 11.42 Hz (−21.82 – 21.89), n = 11, p = 0.64, Wilcoxon signed-rank test; Fig. 4K). Interestingly, these results suggest that fast-spiking interneurons can generate gamma-frequency inhibition independently of AMPA-mediated excitatory input. There was, however, considerable variability between experiments (see Fig. 4K). In some cases, the frequency changed very little, implying an ING mechanism (e.g., Fig. 4F-I). In others, the frequency increased substantially or decreased slightly after blocking AMPA receptors. The mEC apparently wires itself up to produce fast gamma oscillations when presented with theta drive, but this can be achieved in multiple ways. Neurons have homeostatic mechanisms that allow them to produce their characteristic electrical activity resulting in a large variability (2-5 fold) in the specific values for each conductance in different neurons of the same type (Goaillard and Marder, 2021). This phenomenon is called degeneracy (Golowasch et al., 2002) and also applies to circuits. We use computational modeling below to account for some of this variability.

### PV+ interneurons generate fast gamma frequency inhibition in the mEC

Because our findings showed that fast gamma-frequency inhibition persisted even in the absence of AMPA-mediated synaptic input, we sought to determine whether driving PV+ fast-spiking interneurons alone could generate fast gamma-frequency inhibition and whether their activity differed from broader network-driven inhibition. To investigate this, we crossbred PV-Cre and LoxP-ChR2-EYFP transgenic mouse lines to express ChR2 in PV+ interneurons (PV-ChR2). Whole-cell recordings were used to capture voltage activity and synaptic currents during theta frequency optogenetic stimulation of PV+ interneurons.

We found that PV+ fast-spiking interneurons could fire at fast gamma frequencies during theta-frequency optogenetic stimulation (Fig. 5A, B). Some fast-spiking interneurons fired during optogenetic stimulation (50%, n = 2/4) with an average of 4.34 ± 1.83 spikes per theta cycle for active interneurons. The distribution of theta phase spiking was broadly tuned over the center half of the stimulation period (Fig. 5C), as with Thy1+ stimulation. The interspike firing rate of PV+ fast spiking interneurons peaked at nearly 150 Hz (Fig. 5D), exceeding the peak firing frequency observed during Thy1+ stimulation.

**Figure 5:**
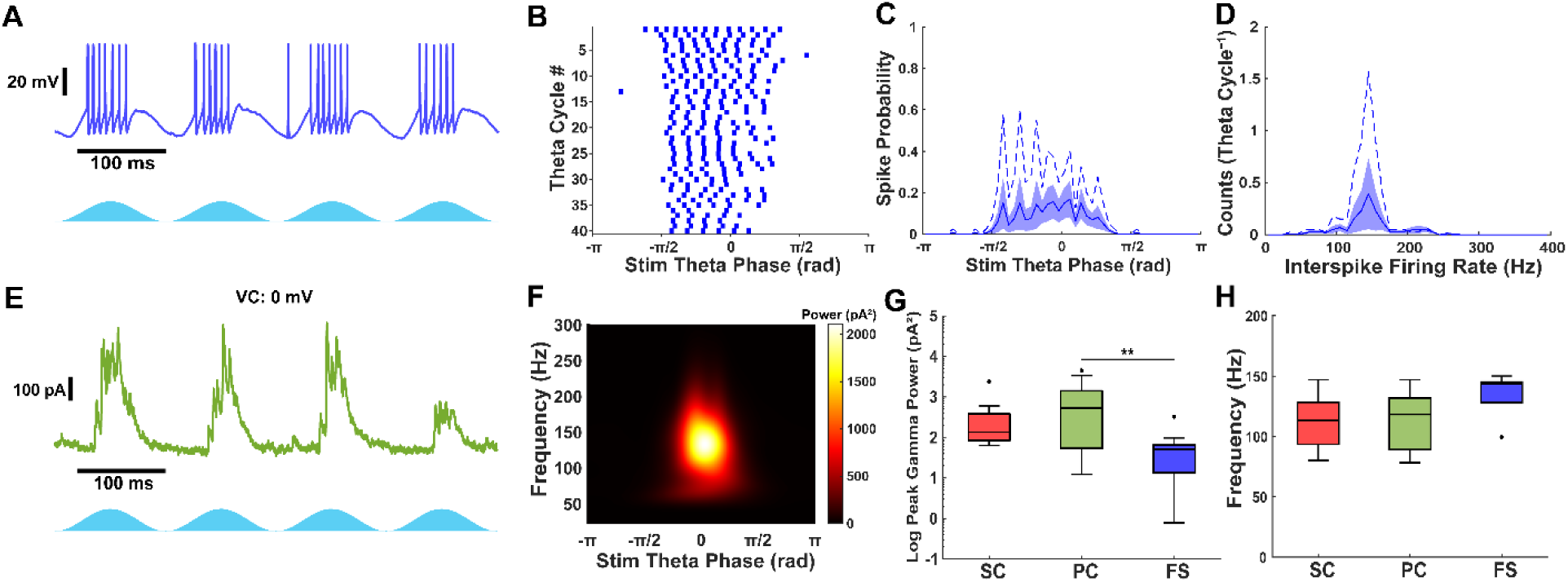
PV+ inhibitory current oscillations are weaker and faster than Thy1. A) Example PV+ interneuron voltage trace during optogenetic stimulation. B) Raster plot of 40 theta cycles from voltage trace in A. C) Theta stimulation spike-phase histogram of fast-spiking interneurons. Solid line and shaded region show mean ± S.E.M. Dashed line represents example shown in A, B. D) Interspike firing rate histogram of fast-spiking interneurons. Solid line and shaded region show mean ± S.E.M. Dashed line represents example shown in A, B. E) Example pyramidal cell inhibitory current trace during theta-frequency PV+ stimulation. F) Average scalogram from 40 theta cycles of inhibitory currents from trace in F. G) PV+ inhibitory current peak gamma power in all cell types. Inhibition is stronger in excitatory cells. H) PV+ inhibitory current gamma frequency. PV+ inhibition (ING) is faster than Thy1+ (PING). Outliers are indicated by filled circles.

All cell types received fast gamma frequency PV+ inhibition during optogenetic stimulation (Fig. 5E, F, H), including PV+ interneurons (Fig. 5-1). Cell type had a significant effect on the peak gamma power of PV+ inhibitory currents (χ^2^(2,41) = 10.92, p = 0.0043, Kruskal–Wallis test). Pyramidal cells received the strongest gamma frequency inhibition (Pyramidal: 531 pA^2^ (55 – 1429), n = 23), significantly greater than fast-spiking interneurons (Fast-spiking: 51 pA^2^ (14 – 66), n = 12; p = 0.0039; Fig. 5G), but not stellate cells (Stellate: 135 pA^2^ (84 – 385), n = 9; p = 1). However, the power of gamma frequency PV+ inhibition was not statistically different between stellate cells and fast-spiking interneurons (p = 0.055), in contrast to what was observed during Thy1+ stimulation. The median peak inhibitory gamma power in stellate cells and fast-spiking interneurons, but not pyramidal cells, was greater during Thy1+ stimulation (Stellate: p = 0.018; Pyramidal: p = 0.76; Fast-spiking: p = 0.011; Mann-Whitney U test), indicating that PV+ interneurons generate weaker inhibitory oscillations compared to network drive of Thy1+ excitatory and inhibitory interneurons.

All cell types received inhibition at similar gamma frequencies during PV+ stimulation (Stellate: 113.2 Hz (93.4 – 128.3), n = 9/9; Pyramidal: 118.2 Hz (89.2 – 131.7), n = 22/23; Fast-spiking: 143.6 Hz (128.0 – 145.2), n = 5/12; χ^2^(2,33) = 5.14, p = 0.077, Kruskal–Wallis test). The median frequency of inhibitory currents was faster during PV+ stimulation in pyramidal cells and fast-spiking interneurons, but not stellate cells compared to Thy1+ stimulation (Stellate: p = 0.11; Pyramidal: p = 1.1e−4; Fast-spiking: p = 0.014; Mann-Whitney U test), indicating that PV+ interneurons synchronize at a slightly faster rate independently.

In one example, we show that optogenetic simulation of PV+ interneurons inhibits the firing of a PV+ cell (Fig. 5-1A, B). Furthermore, this PV+ interneuron receives similar high gamma frequency PV+ inhibition compared to excitatory cells (∼130 Hz; Fig. 5-1C, D). We verified that these currents were GABAergic by washing in 10 µM Gabazine. The blockade of GABA_A_ receptors clearly abolishes gamma frequency currents during optogenetic stimulation (Fig. 5-1E, F). The presence of local PV+ synaptic connectivity is consistent with a previous study that found both synaptic and gap junction connectivity (Fernandez et al., 2022).

These findings demonstrate that PV+ fast-spiking interneurons can independently generate gamma-frequency inhibition and play a central role in maintaining gamma oscillations in the mEC. Furthermore, the faster, weaker inhibition generated during PV+ stimulation suggests a specialized role for PV+ interneurons in modulating local network dynamics to support spatial and memory-related computations.

### Computational modeling of Thy1+ network stimulation in layer II/III mEC

There are two prevailing models (Bartos et al., 2007) for the generation of gamma oscillations, interneuronal network gamma (ING) and pyramidal interneuronal network gamma (PING). In ING, the oscillations are generated by interactions between the interneurons (Wang and Buzsáki, 1996). There is a stochastic ING mechanism in which individual interneurons are subthreshold and fluctuation-driven (Brunel and Hansel, 2006). However, we will not address this mechanism here since the interneurons are suprathreshold due to the theta drive. In PING, the interneurons are generally assumed to be quiescent (Börgers and Kopell, 2003) but are driven to fire by a volley of firing in the excitatory neuron population when that population recovers from the previous wave of inhibition. Although there are two types of excitatory cells in the mEC, stellate and pyramidal cells, for the purpose of our computational model, we will simply call the excitatory cells (‘E cells’), and the inhibitory PV+ interneurons (‘I cells’). We added 400 excitatory (E) stellate cells to our previously calibrated heterogeneous network of 100 PV+ fast-spiking interneurons (Via et al., 2022) and did not incorporate a separate model of the pyramidal cell population. The stellate cells were not connected to each other (Dhillon and Jones, 2000; Couey et al., 2013; Pastoll et al., 2013; Fuchs et al., 2016).

Our computational simulations focused on the relative roles of excitation and inhibition in generating theta-nested gamma in mEC. Previously, optogenetically-driven theta-nested gamma oscillations (∼100 Hz) in mEC slices from Thy1-ChR2 mice were abolished after blocking AMPA synaptic currents (Pastoll et al., 2013), in contrast to our results showing a minimal reduction. Interestingly, Fig. S3D of the same study (Pastoll et al., 2013) clearly shows that blocking excitation reduced but did not abolish interneuronal firing, contrary to classic models of driven I cell PING (Börgers and Kopell, 2003) that require the interneurons to be silent in the absence of synaptic excitation, and as parameterized in their model of gamma generation (Fig. 6-1). Therefore, in our model, the interneurons receive sufficient simulated optogenetic theta drive to fire in the absence of synaptic excitation.

**Figure 6:**
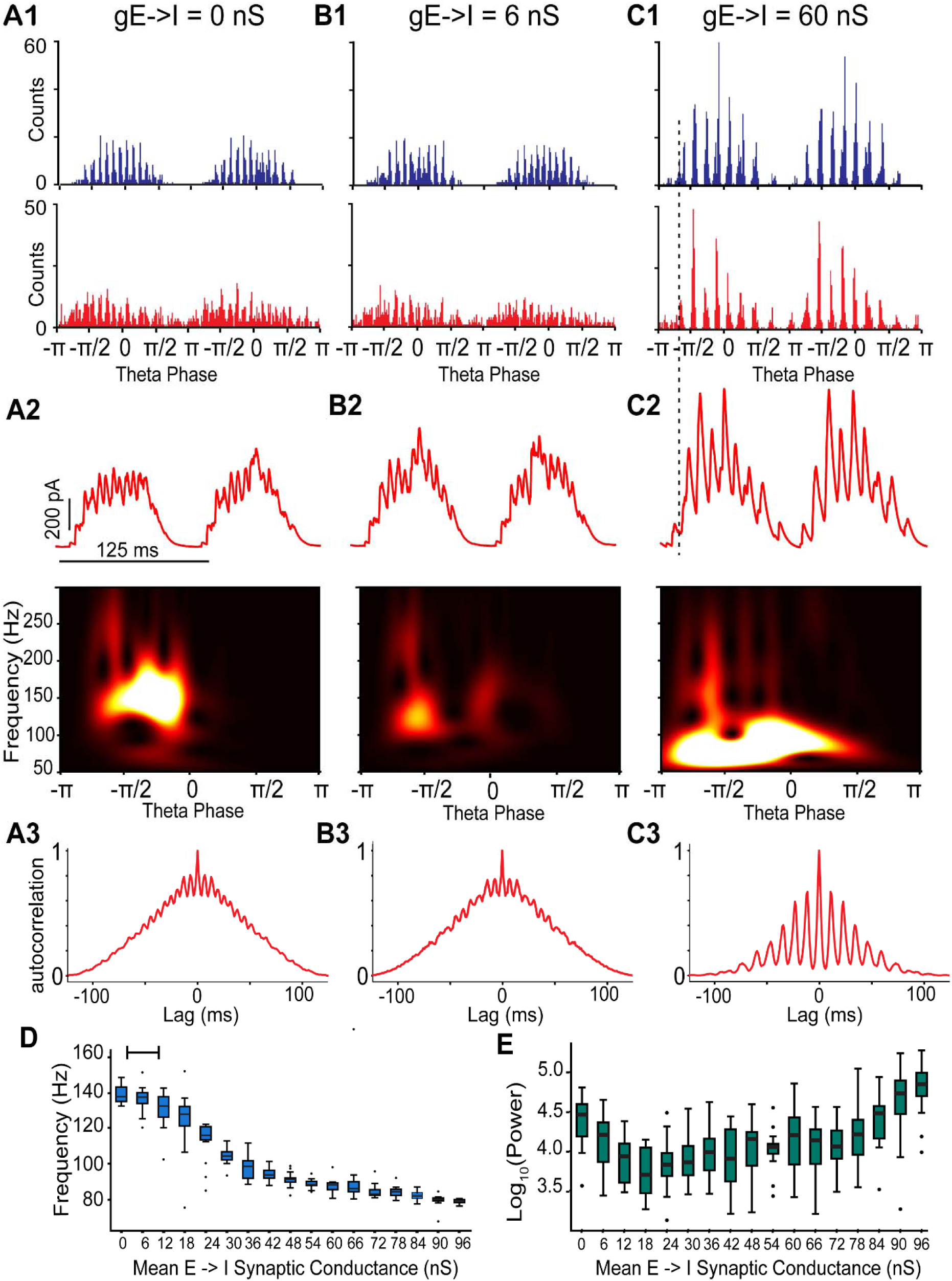
Simulation of excitatory-inhibitory networks with suprathreshold interneuron drive and recurrent inhibition: transitions from ING to PING dominated frequency with increasing excitatory connectivity strength. A) E-I-I network with no E to I conductance to simulate AMPA receptor blockade. B) E-I-I network with weak E to I conductance (6 nS) C) E-I-I network with strong E to I conductance (60 nS). A1-C1: Spike time histograms for I cells (top, blue) and E cells (bottom, red). A2-C2 top: Inhibitory synaptic currents (red) from a representative E cell during simulated optogenetic theta drive. Bottom: Average scalogram of inhibitory synaptic currents from top. A3-C3: Autocorrelogram of spike time histograms in E cells. D, E) Simulations were run on fifteen networks each with a different seed to randomize connectivity. Open circles indicate outliers. D) The peak frequency transitions from a fast ING dominant rhythm to a slower PING dominant rhythm as the E to I synaptic conductance is increased. The bar at the top reflects the approximate range of this conductance in our slice experiments, based on the amplitude of the oscillations in EPSCs in voltage-clamped readout stellate cells. E) The power at the peak frequency increases and plateaus as the E to I synaptic conductance is increased. The connectivity between I to I, E to I and I to E cells was random with fixed probabilities (see Methods).

**Figure 7:**
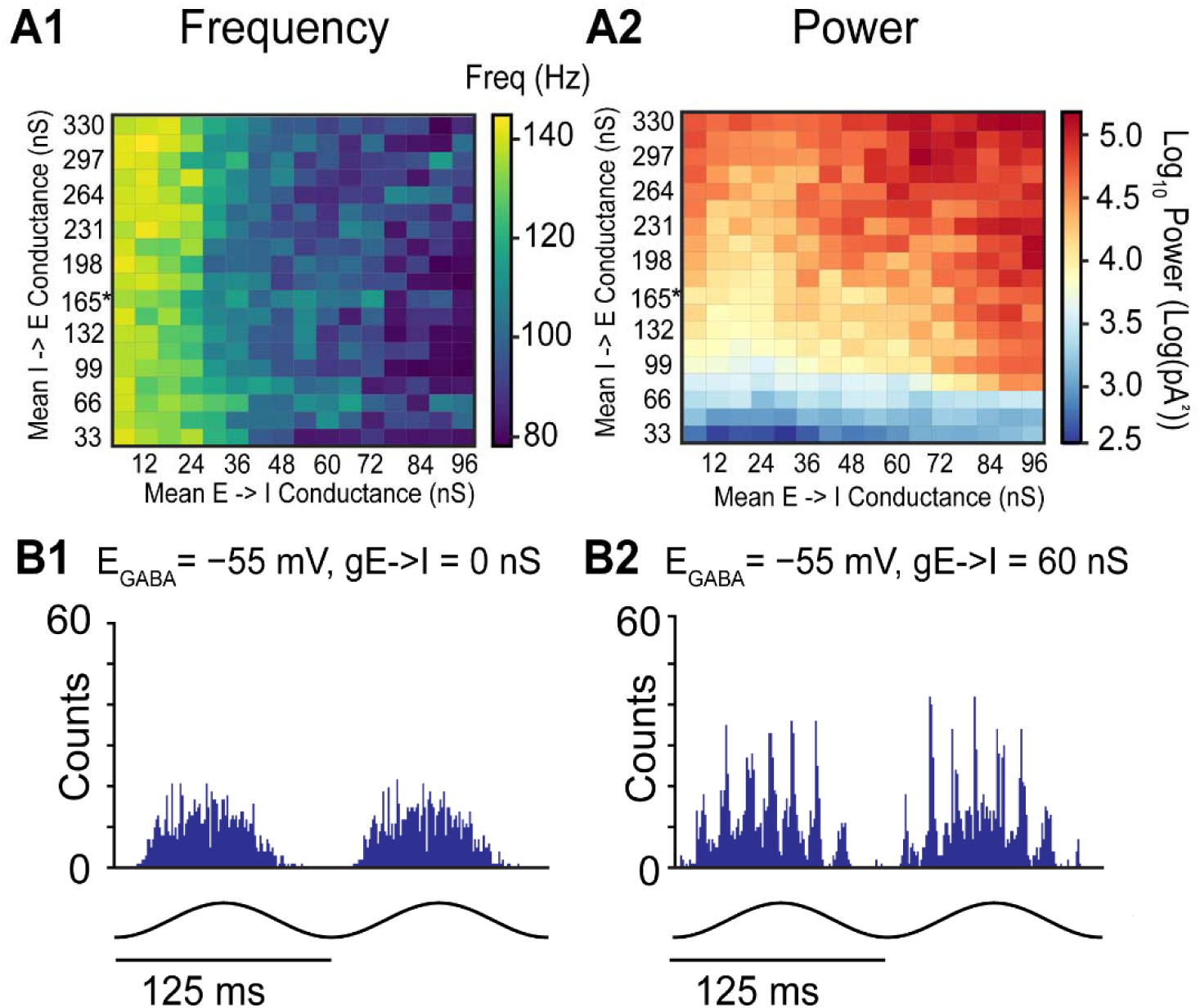
Simulated effects of different I to E connectivity strengths and shunting inhibition. A) Frequency (1) and power (2) heat maps of theta-nested gamma inhibitory currents as I->E and E->I strength is varied. B) Inhibitory cell spike time histograms during simulations with shunting I to I connectivity (E_GABA_ = −55 mV). B1. No E->I connectivity. B2. Strong E->I conductance (60 nS).

During Thy1+ optogenetic stimulation, fast-spiking interneurons receive both theta frequency ChR2 currents and gamma frequency AMPA-mediated synaptic currents when voltage-clamped at −70 mV. To estimate the physiological range of E to I connections, we measured the peak-to-peak amplitude of the gamma frequency excitatory currents from 5 voltage-clamped fast-spiking interneurons (e.g., Fig. 2A bottom, Fig. 4A). These currents were divided by the holding potential (−70 mV) to obtain a range of physiological E to I maximal synaptic conductances (5-10 nS). To replicate the experimental data in this study, hyperpolarizing GABA_A_ synapses (E_GABA_ = −75 mV) were used between interneurons, similar to a previous model (Pastoll et al., 2013; Solanka et al., 2015).

### E to I synaptic conductance strength governs gamma frequency and mechanism

We first show the ability of the network to generate pure ING in response to simulated 8 Hz optogenetic drive (Fig. 6A), analogous to the persistence of gamma (albeit at lower power) after AMPA/kainate receptor blockade in optogenetically stimulated Thy1 mouse slices (Fig. 4H-K) or optogenetic drive of only PV+ interneurons (Fig. 5). Fast gamma modulation is clearly visible in the spike time histograms of both the I cell (Fig. 6A1 top) and the E cell (Fig. 6A1 bottom) populations as well as in the IPSCs received by a model E cell voltage-clamped at 0 mV (Fig. 6A2 top). The peak wavelet frequency (147 Hz, Fig. 6A2 bottom) of the IPSCs received by the E cells corresponds to the reciprocal of the time lag between peaks of the autocorrelogram (143 Hz, Fig. 6A3) of the spike time histogram for the E cells. Importantly, the fast gamma rhythmicity in the E cell population histogram suggests that the ING rhythm is capable of modulating the fast gamma signal transmitted by mEC projection neurons to downstream hippocampal targets such as granule cells in the dentate gyrus or the CA3 or CA1 pyramidal cells. Next, we illustrate the effect of adding E->I connectivity. Weak E to I connections at 6 nS (Fig. 6B) reduced the frequency by ∼10Hz but did not drastically change the frequency of the ING rhythm. Although the power decreased, possibly due to the large amount of noise that was added to prevent the theta drive alone from synchronizing the E cell population. Further strengthening the E->I conductance (60 nS) caused a transition to a slower PING dominated rhythm (Fig. 6C). As indicated by the dashed vertical line (Fig. 6C1, 6C2), once there is sufficiently large E->I conductance, the first population peak in the histogram of the E cell firing (Fig. 6C1 bottom) recruits sufficiently strong I cell firing (Fig. 6C1 top) to prevent the I cells from firing again before the E cells (i.e., E cell recovers first PING). At strong E->I conductances, the firing responses of I cells are grouped into bursts where the PING dominated network frequency is set by the interburst interval, rather than interspike interval of I cells in the ING regime (Fig. 6-2), as observed in Fig. S3D of Pastoll et al. (2013) and occasionally in our study.

Next, we show that increasing the strength of the average total E->I conductance slows the frequency of the theta-nested oscillations (Fig. 6D, 6E), initially by recruiting more inhibition between I cells, but eventually by switching from a driven I cell mechanism to an E cells recover first PING mechanism. Thus either a faster ING dominated or a slower PING dominated oscillation is manifested, although both oscillations are within the fast gamma regime at 65-140 Hz observed *in vivo* in the mEC (Colgin et al., 2009). This mechanism explains much of the variability seen in Fig. 4K, aside from decreases in frequency when DNQX is applied that may reflect experimental variability, or the influence of interneuron subtypes not included in the model (e.g., disinhibition of PV+ cells). When simulations are run with different seeds, each seed has fixed but different connectivity and network parameter distribution. However, in each fixed seed, the conductance noise added to the E cells has a random distribution. This allows us to replicate some of the variability observed in the experimental results.

### Influence of E–I balance and GABA_A_ reversal potentials on gamma oscillations

In prior simulations (Fig. 6), the I->E conductance strength was unchanged. To verify if this trend of frequency and power change was robust to different network connectivity and levels of inhibition, we explored an additional parameter space to generalize the results: the I->E conductance strength. Since the I->E synaptic conductances were based on a log-normal distribution, we varied the mean of the resultant log-normal distribution. The parameter space for the E->I conductance remained same as prior simulations (Fig. 6D, 6E). Frequency and power heatmaps were generated as we varied I->E conductance strength and E->I conductance strength simultaneously for 9 different network connectivity patterns (Fig. 7A). Our simulations demonstrate that for every tested I->E connection strength, increases in E->I strength decreases the average frequency (Fig. 7A1), like the trend observed previously (Fig. 6D, 6E). As expected, increasing I->E conductance strength increases the power of gamma frequency inhibitory currents for every tested E->I conductance strength (Fig. 7A2).

In our models, we utilized connectivity parameters between I cells previously established in our lab (Fernandez et al., 2022; Via et al., 2022). The reversal potential of the GABA_A_ synapses mediated by chloride ion concentrations, has a large impact on the synchronization tendencies of interneuronal networks (Wang and Buzsáki, 1996; Via et al., 2022; Baravalle and Canavier, 2024). This reversal potential (E_GABA_) has three regimes, which we modify here from the definition given by (Bartos et al., 2007) to use the membrane potential during the interspike interval (V_ISI_) rather than the resting membrane potential: 1) a hyperpolarizing regime (E_GABA_ < V_ISI_), 2) a shunting regime in which V_ISI_ ≤ E_GABA_ < action potential threshold (V_thres_) and, 3) an excitatory regime (E_GABA_ ≥ V_thres_). Thus, the reversal potential of GABA_A_ is another dimension of the large parameter space of the network model. In dentate gyrus PV+ interneurons, these synapses were shown to be shunting (Vida et al., 2006). However, in CA1 interneurons the GABA_A_ reversal potential varies considerably, ranging from –75 to –55 mV at the soma, with dendrites exhibiting even more hyperpolarized values (Otsu et al., 2020). Previous models (Pastoll et al., 2013; Solanka et al., 2015) and our simulations (Fig. 6, 7A) have used hyperpolarizing reversal potentials for inhibitory synapses (E_GABA_ = −75 mV). Since not all studies have shown that optogenetically evoked theta-nested gamma oscillations persist in the absence of E to I connectivity (Pastoll et al., 2013), we substituted shunting inhibitory synapses (E_GABA_ = −55 mV) only for the I to I connectivity (Fig. 7B), and found that indeed no ING was observed in the absence of E->I connectivity (Fig. 7B1) whereas the PING mechanism was quite robust with E->I connectivity intact (Fig. 7B2). Therefore, variability in GABA_A_ reversal potentials could be another possible explanation for the conflicting measurements of optogenetically evoked theta-nested gamma oscillations during AMPA blockade under different experimental conditions.

## Discussion

Our findings provide novel insights into the mechanisms underlying gamma oscillations in the medial entorhinal cortex (mEC), emphasizing the role of fast-spiking interneurons in generating and maintaining these rhythms. By leveraging optogenetic stimulation, whole-cell recordings, pharmacological manipulations, and computational modeling, we systematically examined how excitatory and inhibitory neuronal populations contribute to gamma oscillation dynamics.

Experimentally, we found minimal recurrent excitation in pyramidal and stellate cells, consistent with previous most connectivity studies (Dhillon and Jones, 2000; Couey et al., 2013; Pastoll et al., 2013; Fuchs et al., 2016), except for one (Winterer et al., 2017). Gamma-frequency inhibition persisted after blocking AMPA receptors with reduced power, supporting contributions from an interneuron network gamma (ING) mechanism, and selective activation of PV+ interneurons confirmed their ability to independently generate fast gamma rhythms. Complementing these results, our computational modeling demonstrated that weak excitatory input sustains fast ING, while stronger excitation promotes a transition to slower pyramidal-interneuron network gamma (PING). This dual mechanism provides a potential basis for the coexistence and dynamic switching of faster and slower gamma oscillations observed in the entorhinal cortex (Colgin et al., 2009).

### Distinct firings patterns of stellate, pyramidal, and fast-spiking interneurons during Thy1+ stimulation

Our results reveal clear distinctions in the firing properties of stellate, pyramidal, and fast-spiking interneurons during theta-nested gamma oscillations. Fast-spiking interneurons exhibited the highest firing rates and were more likely to fire at fast gamma frequencies (∼100 Hz), whereas stellate and pyramidal neurons displayed lower firing rates and engaged in gamma cycle skipping. These findings align with previous studies indicating that fast-spiking interneurons are key pacemakers of gamma oscillations by providing rhythmic inhibition to excitatory cells (Pastoll et al., 2013). This suggests that subsets of excitatory cell populations are active on different gamma cycles but together provide gamma-frequency excitation onto fast-spiking interneurons.

### Recurrent inhibition plays a dominant role in generating gamma oscillations

Voltage-clamp recordings demonstrated that fast-spiking interneurons receive robust AMPA-mediated excitatory currents at gamma frequencies, unlike stellate and pyramidal neurons, which exhibited minimal gamma-frequency excitation. This supports preferential recruitment of fast-spiking interneurons, consistent with dense excitatory input onto PV+ cells (Beed et al., 2013; Couey et al., 2013). The stronger excitatory drive to interneurons suggests that models of grid cells and theta-nested gamma oscillations in mEC should rely solely on excitatory connectivity to interneurons rather than recurrent excitation. Our computational modeling of biophysically calibrated fully heterogeneous interneuronal networks supports this conclusion, since they still produced robust fast gamma oscillations with no excitatory-excitatory connectivity, in contrast to some previous models which suggest that biophysical levels of heterogeneity destroyed the ING mechanism (Wang and Buzsáki, 1996; White et al., 1998). Further, our modeling suggests that the inhibitory-inhibitory connectivity in our experiments was dominated by hyperpolarizing rather than shunting inhibition.

One of the most striking findings in our study is that gamma-frequency inhibitory currents persisted after AMPA receptor blockade, although at a reduced strength. This suggests that tonic excitation of interneurons in the mEC can generate gamma-frequency inhibition without phasic excitatory input, a circuit mechanism that distinguishes mEC from other cortical areas where gamma oscillations typically depend on phasic excitatory drive (Buzsáki and Wang, 2012). This finding aligns with the ING model of gamma generation, where mutual inhibition among fast-spiking interneurons can sustain rhythmic activity without excitatory input. Consistently, our simulations showed that interneuron networks alone can sustain fast gamma rhythms, underscoring the role of PV+ interneurons as core gamma generators.

### PV+ fast-spiking interneurons as autonomous gamma generators

Optogenetic activation of PV+ interneurons confirmed their capacity to generate fast gamma-frequency inhibition independently. PV+ interneurons fired at faster gamma frequencies compared to fast-spiking interneurons during Thy1+ stimulation, and all cell types received fast gamma-frequency inhibition during PV+ stimulation. The inhibitory currents were significantly stronger in pyramidal cells compared to fast-spiking interneurons, indicating that PV+ interneurons provide robust inhibition to excitatory neurons. However, these gamma oscillations were faster and weaker compared to Thy1+ stimulation, suggesting that other interneuron types could be recruited through feedback inhibition. Additionally, we demonstrate that fast-spiking interneurons receive fast synaptic inhibition from PV+ interneurons (Fig. 5, 5-1), aligning with a previous study in our lab that found high levels of both synaptic and electrical coupling between PV+ interneurons (Fernandez et al., 2022), although another study found only gap junctions at distances greater than 100 microns (Huang et al., 2024). These results support the notion that PV+ interneurons play a central role in coordinating local gamma oscillations and maintaining rhythmic inhibitory drive in the mEC (Buetfering et al., 2014). Since theta oscillations in the hippocampal formation are generally thought to be mediated by GABAergic inhibition of PV+ interneurons (Buzsáki, 2002), there is likely a baseline level of excitation of PV+ interneurons *in vivo* during theta-nested gamma oscillations for the inhibitory theta modulation to be effective (Traub et al., 2023). Although this ING mechanism is now technically a PING mechanism (Bartos et al., 2007) because it would disappear if excitation were blocked. The faster oscillations that result when the PV+ interneurons dominate may be necessary for the activity in the upper range of fast gamma (65–140 Hz) in the mEC (Colgin et al., 2009). On the other hand, the excitatory cell dominated mechanisms may be responsible for the lower end of the fast gamma range, which aligns with our simulations (Fig. 6).

### Reconciling divergent results: a distinct PING mechanism, methodological considerations, and shunting inhibition

There are some differences between our modeling results and a previous model (Solanka et al., 2015), as well as differences between our experimental results and those of a previous study (Pastoll et al., 2013). A previous model found that in the presence of independent noise applied to each E cell, increasing E to I connection strength over relatively small values temporarily increased the gamma frequency prior to decreasing at larger E to I conductances, whereas our study shows a monotonic decrease. Both models used hyperpolarizing inhibition between I cells; however, the regions of the parameter spaces explored are not comparable. In our model, the drive to the I cells is suprathreshold to conform to previous experimental observations (Pastoll et al., 2013; Butler et al., 2018) and our studies of optogenetically driven Thy1+ neurons, where I cells continue to fire when excitation is blocked. Whereas in the Pastoll and Solanka models, the I cells do not fire without E cell input. The transition to a lower gamma frequency as the E to I strength is increased only occurs in the parameter regime with suprathreshold drive to the I cells. Further, our E-I-I model shows that increasing the I to E connection strength increases the power of gamma oscillations while maintaining the same frequency transition as E to I connection strength increases (Fig. 7A), whereas the Solanka model predicts a decrease in gamma frequency as I to E conductance increases. Yet another recent model (Traub et al., 2023) of theta-nested gamma in the mEC produced slower oscillations at 30 Hz, but they were not attempting to model optogenetically induced oscillations.

Experimentally, there are several methodological differences between our Thy1 experiments and Pastoll et al. (2013) that may be relevant for the interpretation of the results: 1) mice were overdosed with isoflurane whereas Pastoll et al. euthanized mice with cervical dislocation, 2) sucrose substitution for NaCl in the slicing ACSF is complete whereas Pastoll et al. used partial substitution, 3) the age range of mice used in our study is 2-6 months old whereas Pastoll et al. studied younger mice aged 6-9 weeks, 4) the optogenetic stimulus used in our study returns to baseline between theta cycles, whereas Pastoll et al. added an offset to maintain a depolarized network in addition to the theta frequency stimulus. Importantly, the firing rates of stellate cells in our study were similar to Pastoll et al. (2013) and resting membrane potentials of stellate cells in our study appear similar to those measured in a different study by the same lab (Pastoll et al., 2020). However, the firing rates of fast-spiking interneurons were lower in our study compared to Pastoll et al. (2013). One explanation for the difference in firing rates could be that Pastoll et al. (2013) used stronger optogenetic stimulation. Pastoll et al. (2013) used light intensities up to 22 mW/mm^2^. In our study, we tested light intensities up to 24 mW/mm^2^, however the median peak light intensity used for fast-spiking interneuron recordings was ∼6 mW/mm^2^. Pastoll et al. (2013) do not provide any further detail on the level of optogenetic stimulus across experiments, which makes an accurate comparison difficult. In Pastoll et al. (2013), the light stimulation consists of an offset in addition to theta frequency drive. Therefore, fast-spiking interneurons are suprathreshold over a greater proportion of the theta cycle in their study (Fig. 1I bottom) compared to ours (Fig. 1C3), likely leading to an increase in firing rate.

Pastoll et al. (2013) found that theta-nested gamma oscillations in the mEC were abolished by blocking excitation, whereas Butler et al. (2018) and our study found that gamma oscillations persisted at a reduced strength. Butler et al. (2018) transgenically expressed ChR2 under the CaMKIIα promotor rather than Thy1 as in the other two studies, so the persistence of gamma activity in Butler et al. (2018) might be attributed to CaMKIIα expression in a subset of PV+ cells, which would align their results with ours. One possible explanation for the lack of excitation block resistant gamma in Pastoll et al. (2013) could be that the chemical synaptic connections between inhibitory cells were weaker, or that the reversal potential of GABA_A_ synapses, which significantly impacts the synchronization of interneuronal networks (Wang and Buzsáki, 1996; Via et al., 2022; Baravalle and Canavier, 2024), was different. Pastoll et al. (2013) recorded inhibitory currents using a holding potential of −50 mV and a chloride reversal potential of −60 mV. Here, we recorded inhibitory currents using a holding potential of 0 mV and a chloride reversal potential of −75 mV. Therefore, it is possible that we could capture weaker gamma rhythmic inhibitory currents in our study. The extracellular solutions used in Pastoll et al. (2013) and our study had similar concentrations of chloride and bicarbonate ions. However, the intracellular concentration of these two ions in the unmonitored neurons which generate the gamma oscillations in slice preparations are unknown. Higher concentrations of intracellular chloride or bicarbonate (Farrant and Kaila, 2007) could have rendered GABA_A_ synapses shunting, accounting for the stronger effect of AMPA block (Fig. 7B).

Our simulations of a shunting network (Fig. 7B) suggest a form of PING in which interneurons fire asynchronously without synaptic excitation but are forced to synchronize when excitatory input is active. The theoretical basis for this form of PING (Chandrasekaran et al., 2011) was recently extended to account for synaptic delays (Vedururu Srinivas and Canavier, 2024). The principle is that the E–I network can be reduced to a 2D discrete map based on slopes of the phase resetting curves (PRC). In a shunting network that cannot synchronize itself, perturbations grow because they are multiplied by a scaling factor greater than one. The slope of the PRC is stabilized by the input from the excitatory cell and enforces synchrony. This E cells recover first PING mechanism does not require inhibitory cells to be quiescent in the absence of synaptic excitation, in contrast to the classic driven I cell PING (Börgers and Kopell, 2003).

### Implications for grid cell function and spatial computation

Our findings have important implications for understanding how grid cells process spatial information. The persistence of gamma oscillations despite blocking fast excitatory input suggests that inhibitory circuits are crucial for maintaining grid cell dynamics. Given that grid cells rely on precise timing of excitatory and inhibitory inputs, our data suggest that PV+ interneurons may regulate spatial coding by imposing temporal constraints on excitatory neuron firing. Many computational models of grid cell activity and theta-gamma coupling have focused on reciprocal excitatory-inhibitory circuits (Pastoll et al., 2013; Widloski and Fiete, 2014). Our results support the addition of inhibitory-inhibitory connectivity to these models, which has been shown to enhance grid stability and increase gamma frequency (Solanka et al., 2015).

The two major models of grid cell activity are single bump and multi-bump models (Shipston-Sharman et al., 2016). Single bump models require synaptic profiles with surround connectivity strongest at about half the sheet width, whereas multi-bump models rely on shorter-distance connections. According to Figure 74 in (Paxinos and Franklin, 2001), the lateral mEC is about 0.8 by 2 mm at its largest extent. In rats, grid modules extended across mediolateral band for distances greater than 1 mm (Stensola et al., 2012). Based on our previous results (Fernandez et al., 2022), chemical synapses between PV+ interneurons, between SST+ interneurons, and chemical synapses from PV+ and SST+ onto excitatory neurons are minimal at distances greater than 100-200 μm. Moreover, our results show that stellate and pyramidal cells receive minimal gamma frequency excitation (Fig. 2), arguing against recurrent synaptic excitation. Therefore, the only possible source of long-range connectivity to support single bump models is from excitatory to inhibitory cells, which has not yet been extensively characterized.

### Conclusion and future directions

In summary, our study provides evidence that fast-spiking interneurons are essential for gamma oscillation generation in the mEC. While AMPA-mediated excitation plays a role in driving interneuron activity, gamma-frequency inhibition persists independently of excitatory input, suggesting a dominant role for inhibitory circuits. PV+ interneurons serve as generators of gamma rhythms, orchestrating network-wide inhibitory dynamics. Complementing these findings, computational modeling revealed that the strength of excitatory input tunes the balance between ING- and PING-dominated regimes, offering a mechanistic explanation for the coexistence and potential switching between faster (100-140 Hz) and slower (60-100 Hz) gamma oscillations. These findings refine existing models of mEC function and emphasize the importance of inhibitory networks in spatial computation and memory processing.

Further studies should explore how gamma oscillations in the mEC interact with other brain regions involved in spatial navigation, such as the hippocampus. Additionally, computational modeling approaches could help elucidate the precise contribution of inhibitory networks to spatial coding. For example, our modeling studies predict that AMPA receptor positive allosteric modulators—which only potentiate the receptors in the presence of glutamate and have been shown to improve performance on cognitive spatial tasks (Suzuki et al., 2021; Radin et al., 2025)—would increase the power and slow the frequency of theta-nested gamma oscillations in the mEC. Investigating gamma oscillation disruptions in neurodegenerative diseases like Alzheimer’s may also provide valuable insights into the role of inhibitory dysfunction in cognitive decline.

## Acknowledgements

This work was supported by NIH R01 NS 054281 to C.C.C. and J.A.W., NIH F31 NS 134309 to B.W., and NSF Grant No. 2018936 to C.C.C.

---

**Figure 1-1:**
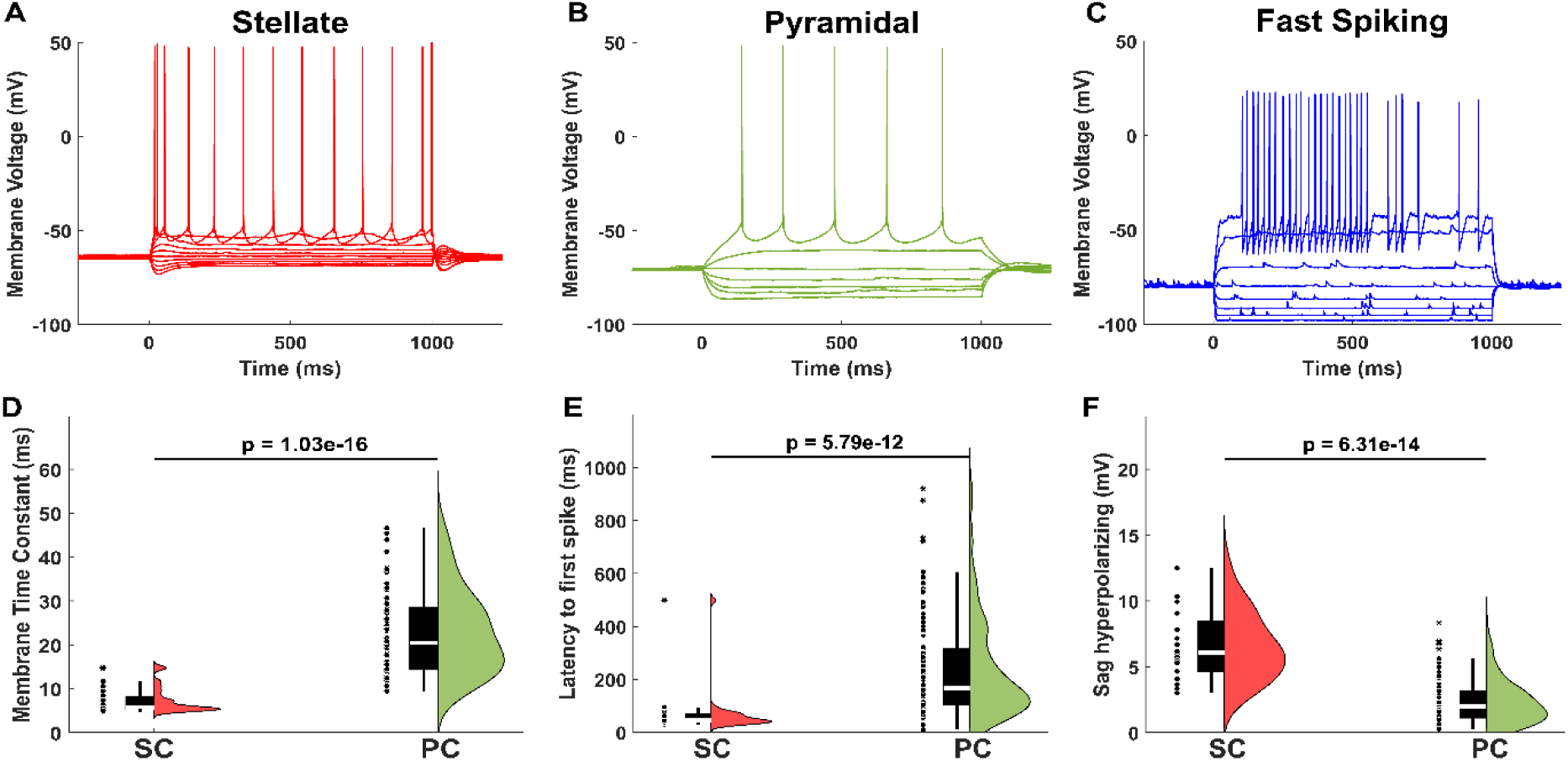
**Classification of major electrophysiological cell types in mEC.** A) Example voltage response to current steps in stellate cell. B) Example voltage response to current steps in pyramidal cell. C) Example voltage response to current steps in fast-spiking interneuron. D) Membrane time constants of stellate and pyramidal cells. Stellate cells have shorter time constants. E) Latency to first spike of stellate and pyramidal cells. Stellate cells fire sooner than pyramidal cells. F) Hyperpolarizing sag potential in stellate and pyramidal cells. Stellate cells have larger hyperpolarizing sag potentials.

**Figure 3-1:**
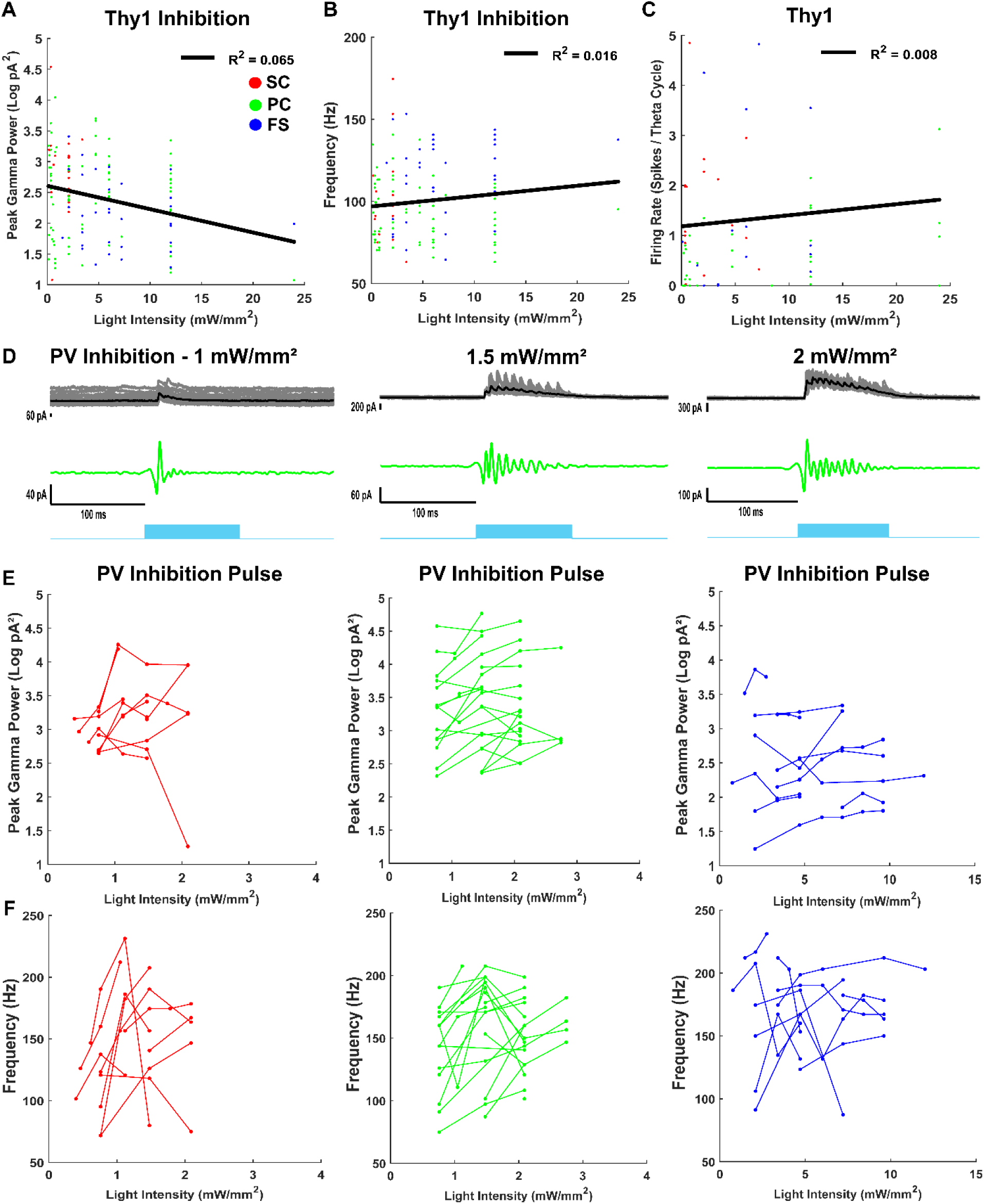
**The effects of optogenetic stimulation intensity on mEC gamma activity.** A) Peak log gamma power of inhibitory currents recorded from stellate cells, pyramidal cells, and fast-spiking interneurons in the mEC vs. peak light intensity of sinusoidal optogenetic Thy1 stimulation. B) Peak gamma frequency of inhibitory currents recorded from stellate cells, pyramidal cells, and fast-spiking interneurons in the mEC vs. peak light intensity of sinusoidal optogenetic Thy1 stimulation. C) Firing rates of stellate cells, pyramidal cells, and fast-spiking interneurons in the mEC vs. peak light intensity of sinusoidal optogenetic Thy1 stimulation. D) Example inhibitory current recordings in pyramidal cell during different levels of pulsed optogenetic PV stimulation (left: 1 mW/mm^2^, middle: 1.5 mW/mm^2^, right: 2 mW/mm^2^). Black line shows average inhibitory currents (20 trials). Gray lines show individual trials. Top traces are raw data. Middle traces are filtered from 50-250 Hz. Light blue square indicates stimulation period. E) Peak gamma power in paired individual cell recordings during different pulse PV stimulation intensities (left: stellate, middle: pyramidal, right: fast-spiking interneuron). Lines indicate paired cell recordings. F) Peak gamma frequency in paired individual cell recordings during different pulse PV stimulation intensities (left: stellate, middle: pyramidal, right: fast-spiking interneuron). Lines indicate paired cell recordings.

**Figure 3-2:**
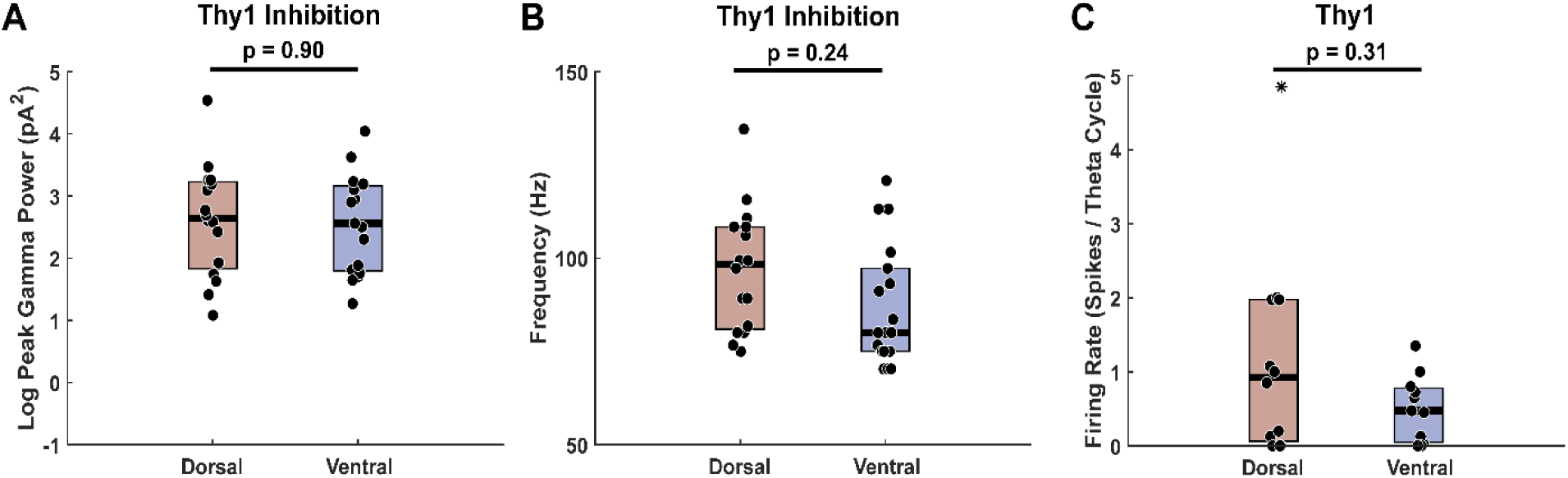
**Gamma frequency inhibition is not different across the dorsal and ventral extents of the mEC during Thy1 stimulation.** A) Peak log gamma power of inhibitory currents recorded from excitatory cells in the most dorsal and ventral mEC slices. B) Peak gamma frequency of inhibitory currents recorded from excitatory cells in the most dorsal and ventral mEC slices. C) Firing rates of excitatory cells in the most dorsal and ventral mEC slices.

**Figure 5-1:**
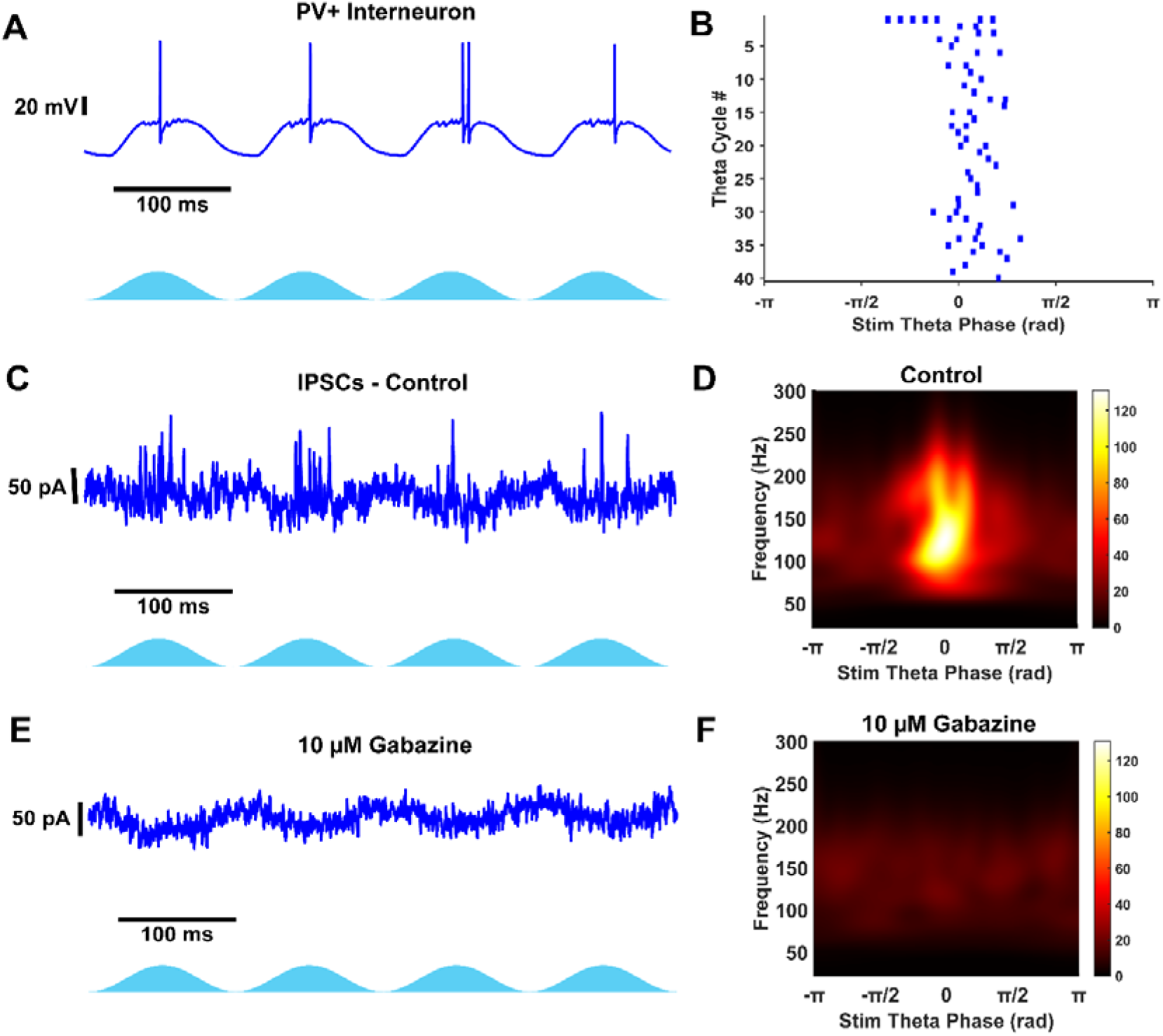
**PV+ interneuron receives fast GABAergic theta-nested gamma inhibition.** A) Example voltage recording in PV+ interneuron during network PV+ optogenetic stimulation. B) Raster plot from 40 theta stimulation periods in same cell as A. C) Voltage clamp at 0 mV records theta-nested IPSCs during PV+ stimulation. D) Average Scalogram from 40 theta stimulation periods of data from example in C. E) Voltage clamp at 0 mV observes no IPSCs after blocking GABA_A_ receptors with 10 µM Gabazine. This data verifies theta-nested gamma in C is fast GABAergic inhibition. F) Average scalogram from 40 theta stimulation periods after blocking GABA_A_ channels. Gamma frequency activity is abolished.

**Figure 6-1:**
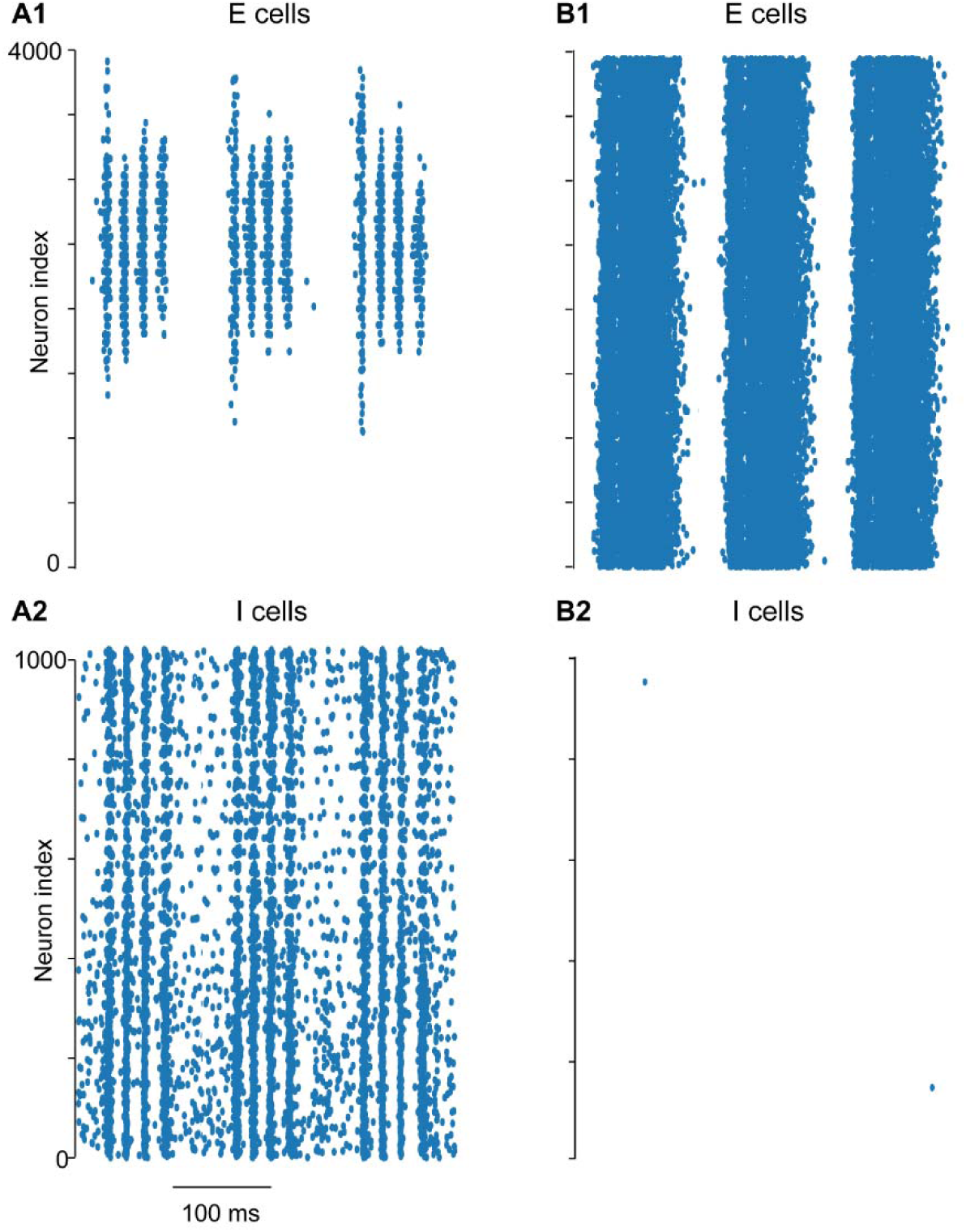
**Simulations of theta-nested gamma oscillations using the grid cell model from Pastoll et al. (2013) with and without AMPA-mediated connections.** A) Raster plots for (1) excitatory cells and (2) inhibitory cells with the default model parameters. B) Raster plots for (1) excitatory cells and (2) inhibitory cells with no AMPA-mediated connectivity. Original model: https://modeldb.science/150031?tab=2&file=GridCellModel/grid_cell_model and a revised simulation_fig_model.py file available at https://github.com/ccanav/pastoll_et_al_2013.

**Figure 6-2:**
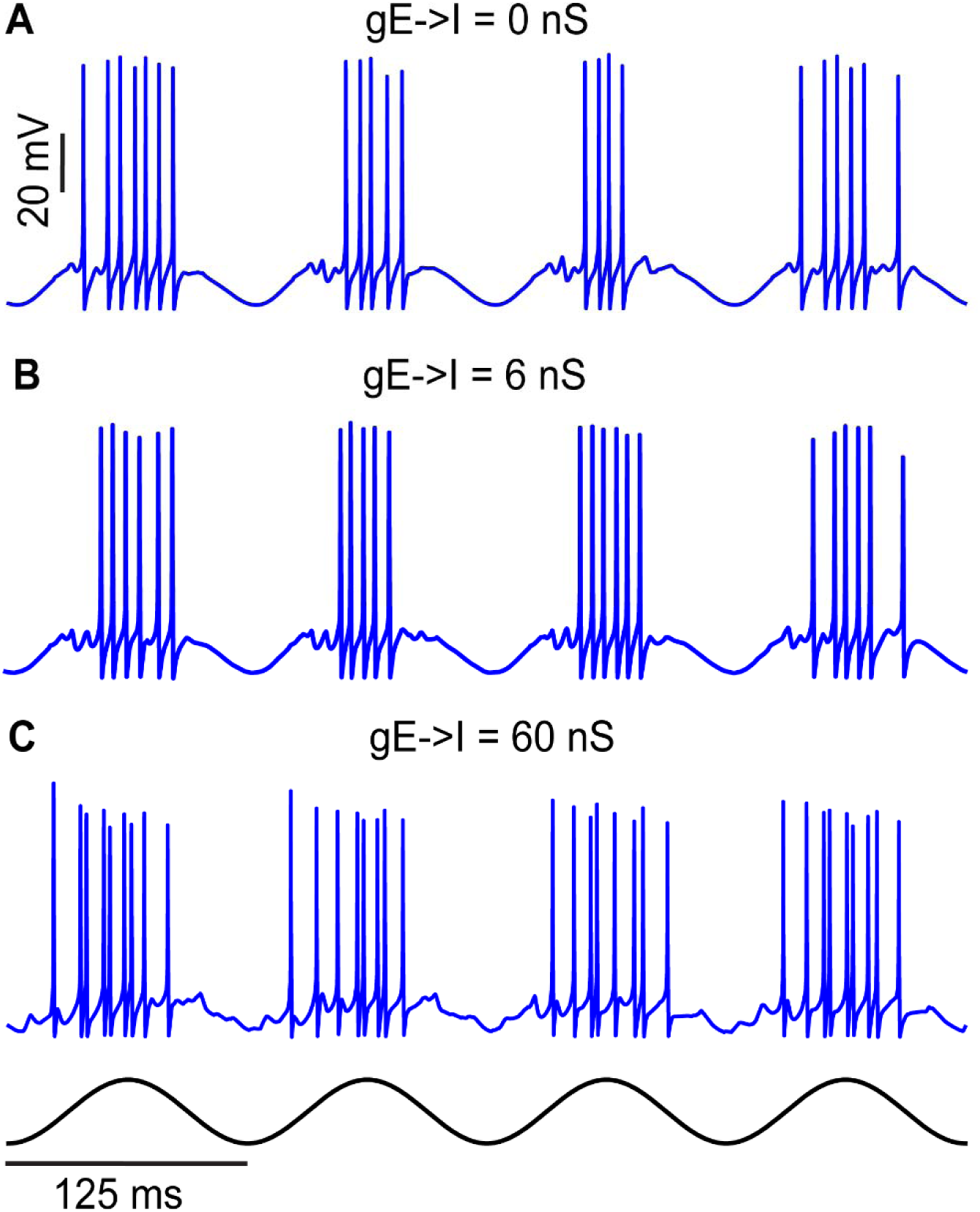
Simulated firing responses of PV interneurons for different E to I connection strengths. A, B) ING regime: With no synaptic excitation (top) and at weak excitation (middle), the I cells fire in a similar pattern. C) PING regime: With strong E to I conductance strength, PV interneuron firing is grouped into bursts, thereby increasing the latency to the next burst of excitation, and slowing the network gamma frequency.

**Extended Data 1.** : The mEC network model and simulation software.

## References

1. Alonso A, Klink R (1993) Differential electroresponsiveness of stellate and pyramidal-like cells of medial entorhinal cortex layer II. Journal of Neurophysiology 70:128–143.

2. Baravalle R, Canavier CC (2024) Synchrony in Networks of Type 2 Interneurons Is More Robust to Noise with Hyperpolarizing Inhibition Compared to Shunting Inhibition in Both the Stochastic Population Oscillator and the Coupled Oscillator Regimes. eNeuro 11:ENEURO.0399-23.2024.

3. Bartos M, Vida I, Jonas P (2007) Synaptic mechanisms of synchronized gamma oscillations in inhibitory interneuron networks. Nature Reviews Neuroscience 8:45–56.

4. Beed P, Gundlfinger A, Schneiderbauer S, Song J, Böhm C, Burgalossi A, Brecht M, Vida I, Schmitz D (2013) Inhibitory Gradient along the Dorsoventral Axis in the Medial Entorhinal Cortex. Neuron 79:1197–1207.

5. Börgers C, Kopell N (2003) Synchronization in Networks of Excitatory and Inhibitory Neurons with Sparse, Random Connectivity. Neural Computation 15:509–538.

6. Börgers C, Kopell N (2005) Effects of noisy drive on rhythms in networks of excitatory and inhibitory neurons. Neural Comput 17:557–608.

7. Börgers C, Walker B (2013) Toggling between gamma-frequency activity and suppression of cell assemblies. Front Comput Neurosci 7:33.

8. Brunel N, Hansel D (2006) How Noise Affects the Synchronization Properties of Recurrent Networks of Inhibitory Neurons. Neural Computation 18:1066–1110.

9. Buetfering C, Allen K, Monyer H (2014) Parvalbumin interneurons provide grid cell–driven recurrent inhibition in the medial entorhinal cortex. Nat Neurosci 17:710–718.

10. Butler JL, Hay YA, Paulsen O (2018) Comparison of three gamma oscillations in the mouse entorhinal–hippocampal system. Eur J of Neuroscience 48:2795–2806.

11. Buzsáki G (2002) Theta Oscillations in the Hippocampus. Neuron 33:325–340.

12. Buzsáki G, Draguhn A (2004) Neuronal Oscillations in Cortical Networks. Science 304:1926–1929.

13. Buzsáki G, Moser EI (2013) Memory, navigation and theta rhythm in the hippocampal-entorhinal system. Nat Neurosci 16:130–138.

14. Buzsáki G, Wang X-J (2012) Mechanisms of Gamma Oscillations. Annu Rev Neurosci 35:203–225.

15. Canto CB, Witter MP (2012) Cellular properties of principal neurons in the rat entorhinal cortex. II. The medial entorhinal cortex. Hippocampus 22:1277–1299.

16. Chandrasekaran L, Achuthan S, Canavier CC (2011) Stability of two cluster solutions in pulse coupled networks of neural oscillators. J Comput Neurosci 30:427–445.

17. Chrobak JJ, Buzsáki G (1998) Gamma Oscillations in the Entorhinal Cortex of the Freely Behaving Rat. Journal of Neuroscience 18:388–398.

18. Colgin LL (2013) Mechanisms and Functions of Theta Rhythms. Annu Rev Neurosci 36:295–312.

19. Colgin LL, Denninger T, Fyhn M, Hafting T, Bonnevie T, Jensen O, Moser M-B, Moser EI (2009) Frequency of gamma oscillations routes flow of information in the hippocampus. Nature 462:353–357.

20. Couey JJ, Witoelar A, Zhang S-J, Zheng K, Ye J, Dunn B, Czajkowski R, Moser M-B, Moser EI, Roudi Y, Witter MP (2013) Recurrent inhibitory circuitry as a mechanism for grid formation. Nat Neurosci 16:318–324.

21. Dhillon A, Jones RSG (2000) Laminar differences in recurrent excitatory transmission in the rat entorhinal cortex in vitro. Neuroscience 99:413–422.

22. Domnisoru C, Kinkhabwala AA, Tank DW (2013) Membrane potential dynamics of grid cells. Nature 495:199–204.

23. Dura-Bernal S, Suter BA, Gleeson P, Cantarelli M, Quintana A, Rodriguez F, Kedziora DJ, Chadderdon GL, Kerr CC, Neymotin SA, McDougal RA, Hines M, Shepherd GM, Lytton WW (2019) NetPyNE, a tool for data-driven multiscale modeling of brain circuits. eLife 8:e44494.

24. Farrant M, Kaila K (2007) The cellular, molecular and ionic basis of GABAA receptor signalling. In: Progress in Brain Research, pp 59–87. Elsevier.

25. Fellous J-M, Rudolph M, Destexhe A, Sejnowski TJ (2003) Synaptic background noise controls the input/output characteristics of single cells in an *in vitro* model of *in vivo* activity. Neuroscience 122:811–829.

26. Fernandez FR, Via G, Canavier CC, White JA (2022) Kinetics and Connectivity Properties of Parvalbumin-and Somatostatin-Positive Inhibition in Layer 2/3 Medial Entorhinal Cortex. eNeuro 9.

27. Fries P (2009) Neuronal Gamma-Band Synchronization as a Fundamental Process in Cortical Computation. Annu Rev Neurosci 32:209–224.

28. Fuchs EC, Neitz A, Pinna R, Melzer S, Caputi A, Monyer H (2016) Local and Distant Input Controlling Excitation in Layer II of the Medial Entorhinal Cortex. Neuron 89:194–208.

29. Fuhs MC, Touretzky DS (2006) A Spin Glass Model of Path Integration in Rat Medial Entorhinal Cortex. J Neurosci 26:4266–4276.

30. Fyhn M, Molden S, Witter MP, Moser EI, Moser M-B (2004) Spatial Representation in the Entorhinal Cortex. Science 305:1258–1264.

31. Goaillard J-M, Marder E (2021) Ion Channel Degeneracy, Variability, and Covariation in Neuron and Circuit Resilience. Annu Rev Neurosci 44:335–357.

32. Golowasch J, Goldman MS, Abbott LF, Marder E (2002) Failure of averaging in the construction of a conductance-based neuron model. J Neurophysiol 87:1129–1131.

33. Hafting T, Fyhn M, Bonnevie T, Moser M-B, Moser EI (2008) Hippocampus-independent phase precession in entorhinal grid cells. Nature 453:1248–1252.

34. Hafting T, Fyhn M, Molden S, Moser M-B, Moser EI (2005) Microstructure of a spatial map in the entorhinal cortex. Nature 436:801–806.

35. Hodgkin AL, Huxley AF (1952) A quantitative description of membrane current and its application to conduction and excitation in nerve. The Journal of Physiology 117:500–544.

36. Huang L-W, Garden DL, McClure C, Nolan MF (2024) Synaptic interactions between stellate cells and parvalbumin interneurons in layer 2 of the medial entorhinal cortex are organized at the scale of grid cell clusters. eLife 12:RP92854.

37. McNaughton BL, Battaglia FP, Jensen O, Moser EI, Moser M-B (2006) Path integration and the neural basis of the “cognitive map.” Nat Rev Neurosci 7:663–678.

38. Miao C, Cao Q, Moser M-B, Moser EI (2017) Parvalbumin and Somatostatin Interneurons Control Different Space-Coding Networks in the Medial Entorhinal Cortex. Cell 171:507–521.e17.

39. Miettinen M, Koivisto E, Riekkinen P, Miettinen R (1996) Coexistence of parvalbumin and GABA in nonpyramidal neurons of the rat entorhinal cortex. Brain Research 706:113–122.

40. Otsu Y, Donneger F, Schwartz EJ, Poncer JC (2020) Cation–chloride cotransporters and the polarity of GABA signalling in mouse hippocampal parvalbumin interneurons. The Journal of Physiology 598:1865–1880.

41. Pastoll H, Garden D, Papastathopoulos I, Sürmeli G, Nolan MF (2020) Inter- and intra-animal variation in the integrative properties of stellate cells in the medial entorhinal cortex. eLife 9:e52258.

42. Pastoll H, Solanka L, van Rossum MCW, Nolan MF (2013) Feedback Inhibition Enables Theta-Nested Gamma Oscillations and Grid Firing Fields. Neuron 77:141–154.

43. Paxinos G, Franklin KBJ (2001) The Mouse Brain in Stereotaxic Coordinates, 2nd ed. San Diego: Academic Press.

44. Proskurina EY, Zaitsev AV (2021) Photostimulation activates fast-spiking interneurons and pyramidal cells in the entorhinal cortex of Thy1-ChR2-YFP line 18 mice. Biochemical and Biophysical Research Communications 580:87–92.

45. Radin DP, Cerne R, Smith JL, Witkin JM, Lippa A (2025) Antipsychotic-like pharmacological profile of the low impact ampakine CX691 (farampator): Implications for the use of low impact ampakines in the treatment of schizophrenia. J Psychiatr Res 186:145–153.

46. Sargolini F, Fyhn M, Hafting T, McNaughton BL, Witter MP, Moser M-B, Moser EI (2006) Conjunctive Representation of Position, Direction, and Velocity in Entorhinal Cortex. Science 312:758–762.

47. Sauer J-F, Strüber M, Bartos M (2012) Interneurons Provide Circuit-Specific Depolarization and Hyperpolarization. J Neurosci 32:4224–4229.

48. Shipston-Sharman O, Solanka L, Nolan MF (2016) Continuous attractor network models of grid cell firing based on excitatory–inhibitory interactions. The Journal of Physiology 594:6547–6557.

49. Sohal VS, Zhang F, Yizhar O, Deisseroth K (2009) Parvalbumin neurons and gamma rhythms enhance cortical circuit performance. Nature 459:698–702.

50. Solanka L, van Rossum MC, Nolan MF (2015) Noise promotes independent control of gamma oscillations and grid firing within recurrent attractor networks. eLife 4:e06444.

51. Stensola H, Stensola T, Solstad T, Frøland K, Moser M-B, Moser EI (2012) The entorhinal grid map is discretized. Nature 492:72–78.

52. Sutton NM, Gutiérrez-Guzmán BE, Dannenberg H, Ascoli GA (2024) A Continuous Attractor Model with Realistic Neural and Synaptic Properties Quantitatively Reproduces Grid Cell Physiology. Int J Mol Sci 25:6059.

53. Suzuki A, Kunugi A, Tajima Y, Suzuki N, Suzuki M, Toyofuku M, Kuno H, Sogabe S, Kosugi Y, Awasaki Y, Kaku T, Kimura H (2021) Strictly regulated agonist-dependent activation of AMPA-R is the key characteristic of TAK-653 for robust synaptic responses and cognitive improvement. Sci Rep 11:14532.

54. Tiesinga P, Sejnowski TJ (2009) Cortical enlightenment: are attentional gamma oscillations driven by ING or PING? Neuron 63:727–732.

55. Traub RD, Whittington MA, Cunningham MO (2023) Simulation of oscillatory dynamics induced by an approximation of grid cell output. Rev Neurosci 34:517–532.

56. Traub RD, Whittington MA, Stanford IM, Jefferys JGR (1996) A mechanism for generation of long-range synchronous fast oscillations in the cortex. Nature 383:621–624.

57. Vedururu Srinivas A, Canavier CC (2024) Existence and stability criteria for global synchrony and for synchrony in two alternating clusters of pulse-coupled oscillators updated to include conduction delays. Mathematical Biosciences 378:109335.

58. Via G, Baravalle R, Fernandez FR, White JA, Canavier CC (2022) Interneuronal network model of theta-nested fast oscillations predicts differential effects of heterogeneity, gap junctions and short term depression for hyperpolarizing versus shunting inhibition Rubin J, ed. PLoS Comput Biol 18:e1010094.

59. Vida I, Bartos M, Jonas P (2006) Shunting Inhibition Improves Robustness of Gamma Oscillations in Hippocampal Interneuron Networks by Homogenizing Firing Rates. Neuron 49:107–117.

60. Wang X-J (2010) Neurophysiological and Computational Principles of Cortical Rhythms in Cognition. Physiological Reviews 90:1195–1268.

61. Wang X-J, Buzsáki G (1996) Gamma Oscillation by Synaptic Inhibition in a Hippocampal Interneuronal Network Model. J Neurosci 16:6402–6413.

62. White JA, Chow CC, Ritt J, Soto-Treviño C, Kopell N (1998) Synchronization and oscillatory dynamics in heterogeneous, mutually inhibited neurons. J Comput Neurosci 5:5–16.

63. Whittington MA, Traub RD, Kopell N, Ermentrout B, Buhl EH (2000) Inhibition-based rhythms: experimental and mathematical observations on network dynamics. International Journal of Psychophysiology 38:315–336.

64. Widloski J, Fiete IR (2014) A model of grid cell development through spatial exploration and spike time-dependent plasticity. Neuron 83:481–495.

65. Winterer J, Maier N, Wozny C, Beed P, Breustedt J, Evangelista R, Peng Y, D’Albis T, Kempter R, Schmitz D (2017) Excitatory Microcircuits within Superficial Layers of the Medial Entorhinal Cortex. Cell Rep 19:1110–1116.

66. Wouterlood FG, Härtig W, Brückner G, Witter MP (1995) Parvalbumin-immunoreactive neurons in the entorhinal cortex of the rat: localization, morphology, connectivity and ultrastructure. J Neurocytol 24:135–153.

